# Efferocytosis Remodels Osteoclast Metabolism and Enhances Antibacterial Defense

**DOI:** 10.64898/2026.10.13.759003

**Authors:** Qixiu Yu, Jie Zhang, Ying Qu, Yong Tang, Yuyang Zhang, Yewei Jia, Qijie Dai, Zhoukai Huang, Jie Shen, Nathachit Limjunyawong, Fei Luo, Ce Dou

**Author notes:** These authors contributed equally (Qixiu Yu, Jie Zhang, Ying Qu). Correspondence to: Fei Luo or Ce Dou.

## Abstract

Osteoclasts are metabolically specialized for bone resorption, but how apoptotic-cell uptake shapes their immune function remains unclear. During *Staphylococcus aureus* bone infection, osteoclasts engulfed apoptotic neutrophils. In osteoclast cultures, apoptotic-neutrophil exposure reduced multinucleation and resorption, induced host-defense programs, and enhanced antibacterial activity, whereas macrophages showed no comparable antibacterial response. MERTK was required for efficient osteoclast efferocytosis. *Ctsk*-Cre-mediated *Mertk* deletion increased bacterial burden and bone loss, whereas transplantation of MERTK-competent bone marrow restored osteoclast efferocytosis and reduced infection-associated bone damage. Transcriptomic and extracellular flux analyses revealed enhanced glycolytic activity and altered lactate handling in efferocytic osteoclasts. Lactate supplementation reduced osteoclast formation without reproducing the antibacterial phenotype. Inhibiting lactate production or export attenuated antibacterial activity in efferocytic osteoclasts, whereas blocking lactate uptake enhanced it. These findings identify osteoclast efferocytosis as an immune adaptation that links apoptotic-neutrophil clearance to metabolic remodeling and antibacterial defense in bone.

## Introduction

Effective host defense must eliminate pathogens while limiting tissue damage. During bone infection, persistent bacteria and osteoclast-mediated resorption damage bone, a mineralized tissue that regenerates slowly and provides mechanical support. Antimicrobial defense must therefore be coordinated with the preservation of bone structure. Neutrophils dominate early antimicrobial responses but accumulate and undergo apoptosis as infection progresses. If apoptotic cells are not promptly removed, secondary necrosis amplifies inflammation and tissue injury^1–3^. Osteoclasts respond to inflammatory cues by resorbing bone, placing them at the interface of neutrophil turnover and tissue damage. How apoptotic-neutrophil clearance influences osteoclast function and antibacterial defense remains poorly understood.

Efferocytosis exposes the engulfing cell to a substantial load of lipids, proteins, and nucleotides that must be degraded, redistributed, or exported^4^. The processing of this cargo can alter cell states and inflammatory responses across homeostatic and pathological settings^5^. In macrophages, apoptotic-cell sensing cooperates with IL-4 or IL-13 signaling to induce programs associated with tissue repair and remodeling^6,7^. Metabolic adaptations support immune resolution and continued efferocytosis^8,9^. Cells with distinct metabolic programs and tissue functions may respond differently to the same apoptotic cargo. How efferocytosis affects cells that retain phagocytic capacity but are specialized for other tissue functions remains unclear.

Osteoclasts share a common myeloid developmental origin with macrophages^10,11^, but differentiate into multinucleated cells whose bone-resorptive activity requires coordinated oxidative phosphorylation and glycolysis^12^. Their metabolic demands and tissue functions therefore differ from those of macrophages. Previous studies have observed osteoclasts binding and engulfing apoptotic cells under defined experimental conditions^13^, but the occurrence and consequences of this process during bone infection remain unclear. Bone infection provides a stringent setting in which apoptotic neutrophils, activated osteoclasts, and bacteria coexist^14,15^. This setting allowed us to examine whether osteoclast uptake of apoptotic neutrophils alters metabolic activity and host defense.

Here, we show that osteoclasts efferocytose apoptotic neutrophils during *Staphylococcus aureus* bone infection. In osteoclast cultures, apoptotic-neutrophil exposure reduces multinucleation and resorption, enhances host-defense programs and antibacterial activity, and elicits metabolic changes distinct from those observed in macrophages. MERTK, a member of the TAM (TYRO3, AXL, and MERTK) family of receptor tyrosine kinases^16^, is required for efficient osteoclast uptake of apoptotic neutrophils. *Ctsk*-Cre-mediated *Mertk* deletion increases bacterial burden and bone loss, whereas transplantation of MERTK-competent bone marrow restores osteoclast efferocytosis and protects against infection-associated bone loss. These findings identify osteoclast efferocytosis as an adaptive response to infection that links apoptotic-cell clearance to metabolic remodeling and immune function in bone.

## Results

### Bone infection induces apoptotic neutrophil accumulation and osteoclast efferocytosis

Pathogen invasion is frequently accompanied by extensive host cell death, and efficient clearance of dying cells is essential for maintaining tissue homeostasis and preventing excessive inflammation^1–3^. Using our established *Staphylococcus aureus*-induced bone infection model^14,15^, we first examined neutrophil accumulation and cell death during disease progression (Fig. 1a).

**Fig. 1.**
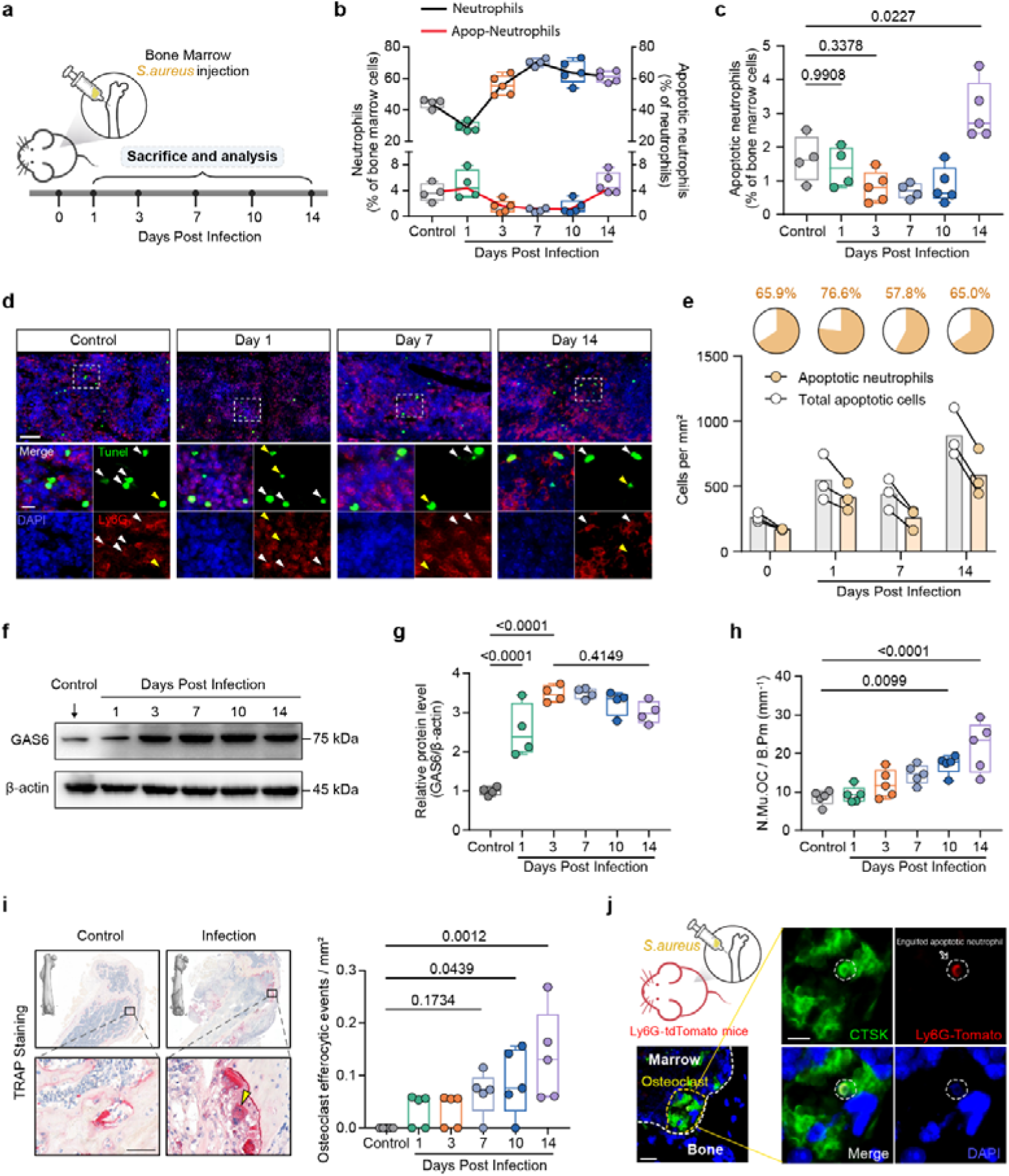
Apoptotic neutrophils accumulate in infected bone and are internalized by osteoclasts. **a**, Schematic of the experimental design for *Staphylococcus aureus*-induced bone infection. **b,c**, Kinetic analysis of bone marrow neutrophil responses after infection, showing neutrophil frequency (**b**) and apoptotic neutrophil frequency (**c**) at the indicated time points. **d,e**, Representative immunofluorescence images of infected bone marrow (**d**) and quantification of apoptotic neutrophils and total apoptotic cells (**e**). Ly6G (red), TUNEL (green), and DAPI staining are shown. White arrows indicate apoptotic neutrophils, and yellow arrows indicate other apoptotic cells. Quantification shows the densities of apoptotic neutrophils and total apoptotic cells (cells/mm²), together with the proportion of apoptotic cells identified as neutrophils. Scale bars, 50 µm (main images) and 10 µm (insets). **f,g**, Immunoblot analysis of GAS6 expression in whole bone marrow lysates collected at the indicated time points after infection (**f**), with densitometric quantification normalized to β-actin (**g**). **h**, Quantification of osteoclast abundance at the indicated time points following bone infection, expressed as N.Mu.OC/B.Pm (number of multinucleated osteoclasts per bone perimeter). **i**, Representative TRAP-stained femoral sections from uninfected and infected mice, with quantification of osteoclasts containing apoptotic cell-associated signals. Arrows indicate osteoclasts undergoing efferocytosis. Scale bars, 50 µm. **j**, Representative bone sections from Ly6G-tdTomato reporter mice after infection showing CTSK⁺ osteoclasts (green), tdTomato⁺ neutrophil-derived signals (red), and DAPI staining (blue). Arrows indicate neutrophil-derived signals within osteoclast boundaries. Scale bars, 20 µm (main images) and 10 µm (insets). Data are presented as mean ± s.d. Statistical significance was assessed using one-way ANOVA with Tukey’s multiple comparisons test (**c,g,h,i**). n = 4 mice per group for control, day 1 and day 7 and n = 5 for day 3, day 10 and day 14 (**b,c**); n = 3 mice per time point (**e**), n = 4 (**g**), and n = 5 (**h,i**). Each point represents one mouse; linked points in e denote measurements from the same mouse.

Flow cytometric analysis revealed changes in the proportions of neutrophils and apoptotic neutrophils following bacterial challenge (Fig. 1b, c and Fig. S1a, b). Consistently, immunofluorescence analysis demonstrated marked accumulation of apoptotic neutrophils, which represented the predominant apoptotic cell population in the infected bone microenvironment (Fig. 1d, e). We next examined whether this accumulation was accompanied by changes in apoptotic-cell recognition machinery. Immunoblotting showed that GAS6, a phosphatidylserine-binding bridging molecule that facilitates apoptotic cell recognition and engulfment^17^, increased after infection and remained elevated (Fig. 1f, g), consistent with previous observations^18^ (Fig. S1c).

As infection progressed, osteoclast abundance increased (Fig. 1h). Histological analysis revealed disrupted bone architecture and apoptotic cell-associated signals within osteoclasts (Fig. 1i), suggesting that osteoclasts participate in apoptotic cell clearance during infection. Apoptotic cell-associated signals within osteoclasts were not detected in uninfected bones but were observed in both the metaphyseal region (Fig. 1i) and the diaphyseal region (Fig. S1d) of infected bones. To test whether osteoclast-associated material originated from neutrophils, we used Ly6G-tdTomato reporter mice to trace neutrophil-derived signals *in vivo*. The presence of tdTomato-positive material within osteoclast boundaries supported the uptake of neutrophil-derived material during bone infection (Fig. 1j and Fig. S1e).

Although efferocytosis is predominantly mediated by professional phagocytes, including macrophages and dendritic cells, specialized nonprofessional phagocytes, such as Sertoli cells and retinal pigment epithelial cells, can also contribute to apoptotic cell clearance and tissue homeostasis^4^. Our findings indicate that osteoclasts also participate in apoptotic-neutrophil clearance during bone infection.

### MERTK mediates apoptotic neutrophil efferocytosis by osteoclasts

Given their shared developmental origin with macrophages^10,11^, we asked whether osteoclasts retain components of the efferocytic machinery during terminal differentiation. The TAM receptor family (TYRO3, AXL, and MERTK) constitutes a major efferocytic receptor system involved in the recognition and uptake of apoptotic neutrophils by phagocytes^7,19^. Reanalysis of published transcriptomic datasets^20,21^ revealed persistent expression of TAM receptor genes during M-CSF- and RANKL-induced osteoclast differentiation in both humans and mice (Fig. 2a and Fig. S2a).

**Fig. 2.**
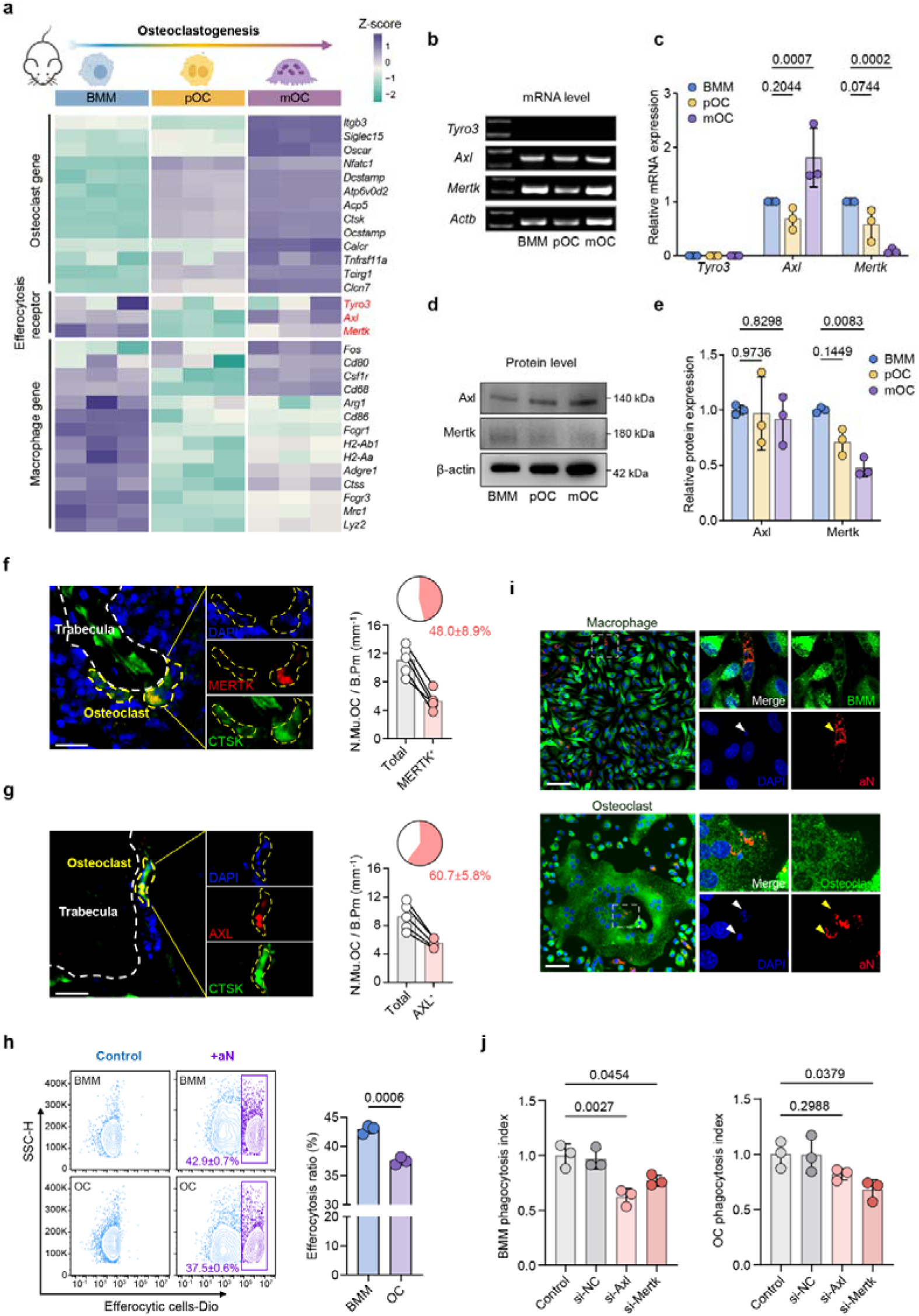
MERTK-dependent efferocytosis of apoptotic neutrophils by osteoclasts. **a**, Heatmap showing expression of osteoclast-associated genes, TAM receptor family genes, and macrophage-associated genes during murine osteoclast differentiation (GSE147174). BMM, bone marrow-derived macrophages (M-CSF, 2 days); pOC, pre-osteoclasts (BMM + RANKL for 1 day); mOC, mature osteoclasts (BMM + RANKL for 3 days). n = 3 bulk RNA-seq samples per differentiation stage; each heatmap column represents one sample. **b**, Representative gel electrophoresis images of PCR products for *Tyro3*, *Axl*, *Mertk*, and *Actb* in BMMs, pOCs, and mOCs. **c**, Quantitative PCR analysis of *Tyro3*, *Axl*, and *Mertk* transcript levels in BMMs, pOCs and mOCs. **d,e**, Immunoblot analysis of AXL, MERTK, and β-actin expression in BMM, pOC, and mOC (**d**), with densitometric quantification (**e**). **f,g**, Immunofluorescence images of mouse femoral sections showing MERTK (**f**) and AXL (**g**) expression in bone-resident osteoclasts. MERTK and AXL are shown in red in **f** and **g**, respectively; CTSK is shown in green, and nuclei are shown in blue (DAPI). Quantification shows the number of receptor-positive multinucleated osteoclasts per bone perimeter and the percentage of osteoclasts positive for MERTK or AXL. Scale bars, 20 µm. **h**, Flow cytometric quantification of apoptotic neutrophil (aN) efferocytosis by BMMs and osteoclasts. **i**, Representative immunofluorescence images of BMMs and osteoclasts incubated with apoptotic neutrophils (aN). Phagocytes are shown in green, apoptotic neutrophils in red, and nuclei stained with DAPI (blue). White arrows indicate apoptotic cells with pyknotic nuclei, and yellow arrows indicate aN. Scale bars, 50 µm. **j**, Quantification of apoptotic neutrophil uptake by BMMs and osteoclasts following siRNA-mediated depletion of *Axl* or *Mertk*. The phagocytosis index was calculated as experimental engulfment (%) divided by control engulfment (%). Data are presented as mean ± s.d. Statistical significance was assessed using two-tailed unpaired *t*-tests (**h**) and one-way ANOVA followed by Tukey’s multiple comparisons test (**c,e,j**). n = 3 independent biological samples per group (**c,e,h,j**) and n = 5 mice (**f,g**). Each point represents one biological sample or one mouse, respectively; linked points in **f,g** denote measurements from the same mouse.

Although *Tyro3* transcripts were undetectable in both macrophages and osteoclasts, *Axl* and *Mertk* expression remained readily detectable by PCR (Fig. 2b, c). Immunoblotting confirmed the presence of AXL and MERTK in differentiated osteoclasts (Fig. 2d, e). MERTK abundance decreased during differentiation but remained detectable in osteoclasts. Consistently, immunofluorescence staining of bone-resident osteoclasts demonstrated AXL and MERTK expression *in vivo* (Fig. 2f, g), supporting a molecular basis for osteoclast-mediated apoptotic cell recognition.

As apoptotic-cell identity shapes the functional responses of efferocytic phagocytes^22^, we next examined whether osteoclasts could directly engulf apoptotic neutrophils, the predominant dying cell population during bone infection (Fig. 1e).

Primary osteoclasts efficiently internalized apoptotic neutrophils, although with slightly lower uptake efficiency than macrophages (Fig. 2h and Fig. S2b). Similarly, RAW264.7 cells and their osteoclast derivatives retained the capacity to engulf apoptotic neutrophils, with lower efficiency than their primary counterparts (Fig. S2c). Because mature multinucleated osteoclasts are poorly captured by cytometry-based assays, we further used confocal microscopy to visualize apoptotic neutrophil signals within osteoclasts, supporting uptake by multinucleated osteoclasts (Fig. 2i).

Given that efferocytic receptor dependency varies among phagocytes and apoptotic cargoes^22–24^, we next investigated the receptor requirements for osteoclast-mediated apoptotic neutrophil clearance. Silencing either *Axl* or *Mertk* impaired apoptotic neutrophil uptake by macrophages. In contrast, osteoclast efferocytosis was reduced by *Mertk* depletion, whereas no significant reduction was detected after *Axl* silencing (Fig. 2j and Fig. S2d). These findings identify MERTK as a key receptor for apoptotic neutrophil uptake by osteoclasts.

### Apoptotic neutrophil efferocytosis remodels osteoclast function and induces host defense programs

Efferocytosis contributes to inflammatory resolution, tissue repair, and immune homeostasis^2,25–28^. We therefore asked how apoptotic-neutrophil exposure affects osteoclast differentiation and function. Apoptotic neutrophils were added during RANKL-induced osteoclast differentiation. At the assay endpoint, aN-treated osteoclast cultures remained TRAP-positive but exhibited reduced multinucleation and a smaller resorbed area compared with controls (Fig. 3a, b).

**Fig. 3.**
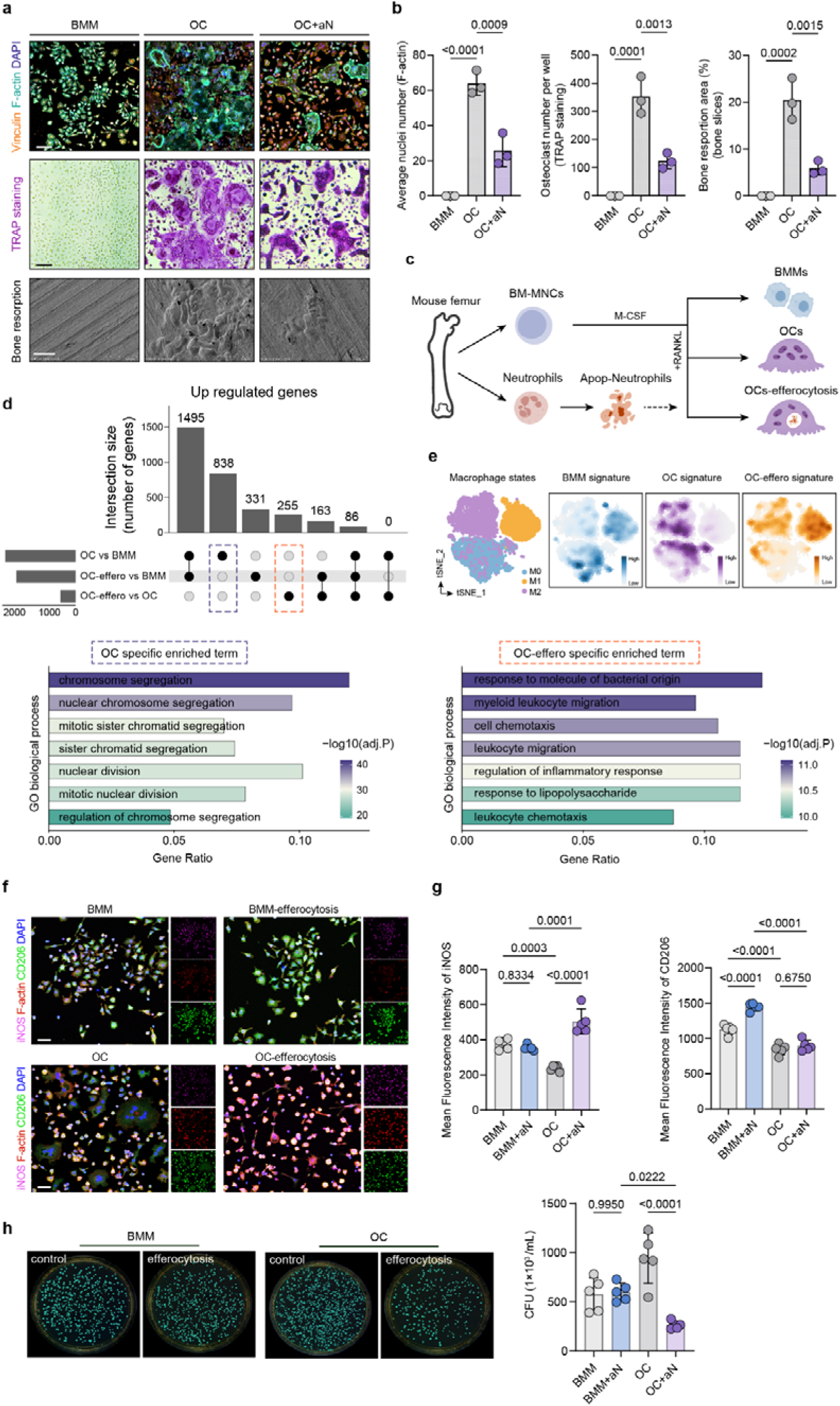
Osteoclast efferocytosis induces host-defense programs and antibacterial activity. **a**, Representative immunofluorescence and TRAP staining images of osteoclasts. F-actin (cyan), vinculin (orange), and DAPI (blue) staining are shown. Scanning electron microscopy images of bone resorption are also shown. Scale bars, 100 µm (immunofluorescence and TRAP images) and 50 µm (scanning electron microscopy images). **b**, Quantification of average nuclear number, TRAP-positive multinucleated osteoclast number per well, and resorbed bone area (%). **c**, Schematic of the experimental design for transcriptomic profiling. BM-MNCs, bone marrow mononuclear cells. **d**, UpSet plot showing the overlap of upregulated genes among the indicated comparisons, together with Gene Ontology biological process enrichment analysis of genes uniquely upregulated during osteoclast differentiation or following apoptotic neutrophil efferocytosis. Analyses used n = 3 biological RNA-seq replicates per group. **e**, t-SNE visualization of macrophage states spanning different polarization programs, showing the enrichment of BMM, osteoclast, and efferocytic osteoclast transcriptional signatures across the macrophage activation landscape. Each point represents one cell from the reference single-cell dataset; these cells are not independent biological replicates. **f,g**, Immunofluorescence analysis of CD206 and iNOS expression in BMMs and osteoclasts before and after apoptotic neutrophil efferocytosis (**f**), with quantification (**g**). CD206 is shown in green, iNOS in magenta, F-actin in red, and nuclei in blue (DAPI). Scale bars, 50 µm. **h**, Viable bacterial counts following challenge with *Staphylococcus aureus*. Representative colony-forming unit (CFU) images and quantification are shown. Data are presented as mean ± s.d. Statistical significance was assessed using one-way ANOVA followed by Tukey’s multiple comparisons test (**b,g,h**). n = 3 independent biological samples per group (**b**) and n = 5 (**g,h**). Each point represents one biological sample.

To characterize these functional changes, we performed bulk transcriptomic profiling of macrophage, osteoclast, and aN-treated osteoclast cultures (Fig. 3c). Genes upregulated during osteoclast differentiation were enriched for biological processes related to mitotic progression and chromosome segregation (Fig. 3d, lower left). Genes upregulated after apoptotic-neutrophil exposure were enriched for responses to bacterial stimuli and innate immune functions (Fig. 3d, lower right).

Consistent with the enrichment results, expression of osteoclast differentiation-associated genes, including *Ctsk* and *Acp5*, was reduced, whereas immune activation-associated genes, including *Cd80*, *Cd86*, and *Cxcl2*, were increased, as confirmed by qPCR (Fig. S3a–c).

To characterize the host-defense program associated with osteoclast efferocytosis, we compared the osteoclast transcriptional profiles with established macrophage polarization signatures^29^. Efferocytic osteoclasts showed greater transcriptional similarity to classical activation and host-defense programs than to reparative macrophage programs (Fig. 3e). MacSpectrum analysis^29^ likewise placed efferocytic osteoclasts toward a host-defense-associated transcriptional state (Fig. S3d).

Immunofluorescence analysis revealed that apoptotic neutrophil uptake increased CD206 expression in macrophages, whereas efferocytic osteoclasts exhibited increased iNOS expression, indicating that the same efferocytic stimulus elicited distinct responses in macrophages and osteoclasts (Fig. 3f, g). Efferocytic osteoclast cultures yielded fewer viable *Staphylococcus aureus* than untreated osteoclast cultures, indicating enhanced antibacterial activity; no comparable reduction was observed in macrophage cultures (Fig. 3h). Given that apoptotic-cell uptake cooperates with IL-4 or IL-13 signaling to induce anti-inflammatory and tissue-repair programs in macrophages^7^, we examined whether cytokine treatment further modified the osteoclast response. Neither TNF-α nor IL-4 produced a comparable enhancement (Fig. S3e, f).

### Efferocytosis rewires osteoclast metabolism toward aerobic glycolysis

Immune effector functions are often accompanied by metabolic adaptations^30–32^. Transcriptomic analysis identified regulators of glucose metabolism alongside immune effector genes among the transcripts strongly upregulated following apoptotic neutrophil uptake (Fig. S4a, b). These observations prompted us to investigate whether efferocytosis induces metabolic reprogramming in osteoclasts.

Consistent with previous reports that osteoclasts are metabolically active cells^12^, comparison of macrophages and osteoclasts revealed higher expression of genes involved in glycolysis, the tricarboxylic acid cycle, and oxidative phosphorylation (OXPHOS) in osteoclast cultures. Apoptotic-neutrophil uptake enhanced glycolytic gene programs while reducing signatures associated with the tricarboxylic acid cycle and OXPHOS (Fig. 4a and Fig. S4c). Immunoblot analysis further confirmed increased expression of GLUT1 and HIF-1α, key regulators of glycolytic adaptation^33–35^, whereas the abundance of mitochondrial OXPHOS proteins decreased following efferocytosis (Fig. 4b). Macrophages did not show comparable changes in these proteins following apoptotic neutrophil uptake.

**Fig. 4.**
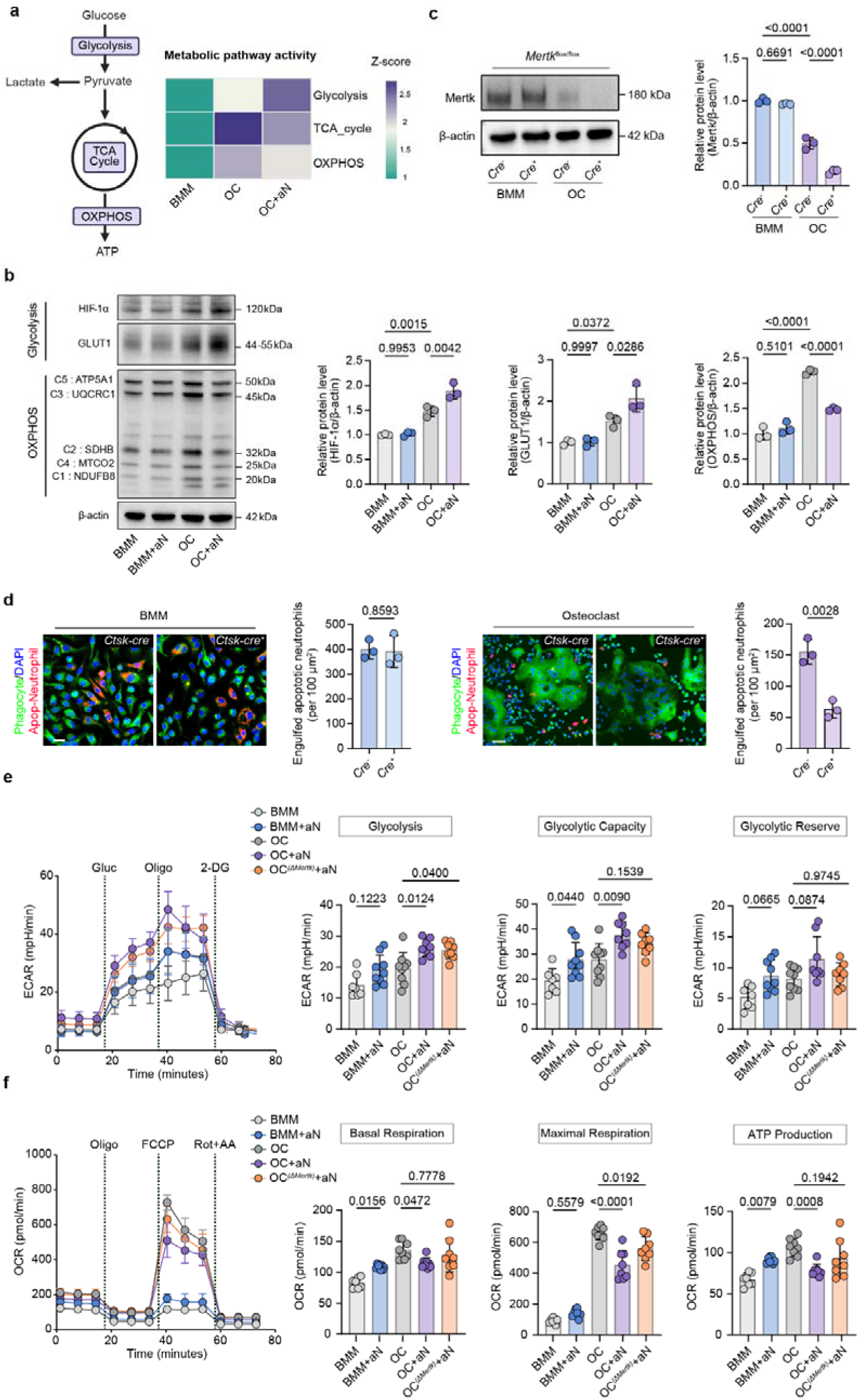
Efferocytosis reprograms energy metabolism in osteoclasts. **a**, Overview of glucose metabolism and associated metabolic pathways, with a heatmap summarizing transcriptional signatures of glycolysis, the tricarboxylic acid cycle, and oxidative phosphorylation across BMMs, osteoclasts, and efferocytic osteoclasts. Bulk RNA-seq analyses used n = 3 biological replicates per group. **b**, Immunoblot analysis of HIF-1α, GLUT1, and proteins detected using an OXPHOS antibody cocktail in BMMs and osteoclasts before and after apoptotic neutrophil efferocytosis, with densitometric quantification. **c**, Immunoblot analysis of MERTK expression in BMMs and osteoclasts derived from OC(ΔMertk) mice and control littermates, with densitometric quantification normalized to β-actin. **d**, Immunofluorescence analysis of apoptotic neutrophil uptake by BMMs and osteoclasts derived from OC(ΔMertk) mice and control littermates. Engulfed apoptotic neutrophils were quantified per 100 µm². Scale bars, 20 µm (BMMs) and 50 µm (osteoclasts). **e,f**, Extracellular flux analysis of glycolytic activity and mitochondrial respiration following apoptotic neutrophil efferocytosis. Glycolytic activity was assessed by extracellular acidification rate (ECAR, mpH/min) (**e**), and mitochondrial respiration was assessed by oxygen consumption rate (OCR, pmol/min) (**f**). The oligomycin-sensitive component of OCR represents ATP-linked respiration. Data are presented as mean ± s.d. Statistical significance was assessed using two-tailed unpaired *t*-tests (**d**) and one-way ANOVA followed by Tukey’s multiple comparisons test (**b,c,e,f**). n = 3 independent biological samples per group (**b–d**); each point represents one biological sample. n = 7–9 measured wells per group (**e,f**); each point in the summary plots represents one well.

To examine the role of MERTK in osteoclast metabolic remodeling, we generated mice with *Ctsk*-Cre-mediated *Mertk* deletion (*Ctsk*-Cre; *Mertk*^flox/flox^), hereafter referred to as OC(ΔMertk) mice. Conditional deletion of *Mertk* (Fig. 4c) selectively impaired apoptotic neutrophil uptake by osteoclasts while preserving macrophage efferocytosis (Fig. 4d). Under uninfected conditions, OC(ΔMertk) mice showed no overt skeletal abnormalities or significant differences in the measured micro-CT parameters (Fig. S4d–f).

We next examined the metabolic consequences of apoptotic neutrophil uptake using Seahorse extracellular flux analysis. Following a 2-h efferocytosis pulse, non-engulfed apoptotic neutrophils were removed, and metabolic activity was assessed after an additional 12 h to capture the sustained effects of efferocytosis. Efferocytosis increased glycolytic capacity in macrophages and both glycolysis and glycolytic capacity in osteoclasts, as assessed by extracellular acidification rate (ECAR) (Fig. 4e). In macrophages, this increase was accompanied by higher basal and ATP-linked respiration, as assessed by oxygen consumption rate (OCR). In contrast, efferocytic osteoclasts displayed increased ECAR together with reduced OCR, consistent with increased glycolytic activity and reduced mitochondrial respiration (Fig. 4e, f). In aN-treated osteoclasts, *Mertk* deletion blunted these changes in ECAR and OCR, indicating that MERTK-mediated efferocytosis rewires osteoclast metabolism toward aerobic glycolysis.

### Osteoclast efferocytosis alters lactate metabolism alongside antibacterial activity

The shift toward aerobic glycolysis in efferocytic osteoclasts (Fig. 4e, f; summarized in Fig. 5a) prompted us to investigate alterations in lactate metabolism. Lactate generated during efferocytosis can regulate phagocyte responses as a signaling metabolite^9,26,36,37^. We therefore investigated whether lactate metabolism contributed to the distinct functional outcomes of efferocytosis in osteoclasts and macrophages.

**Fig. 5.**
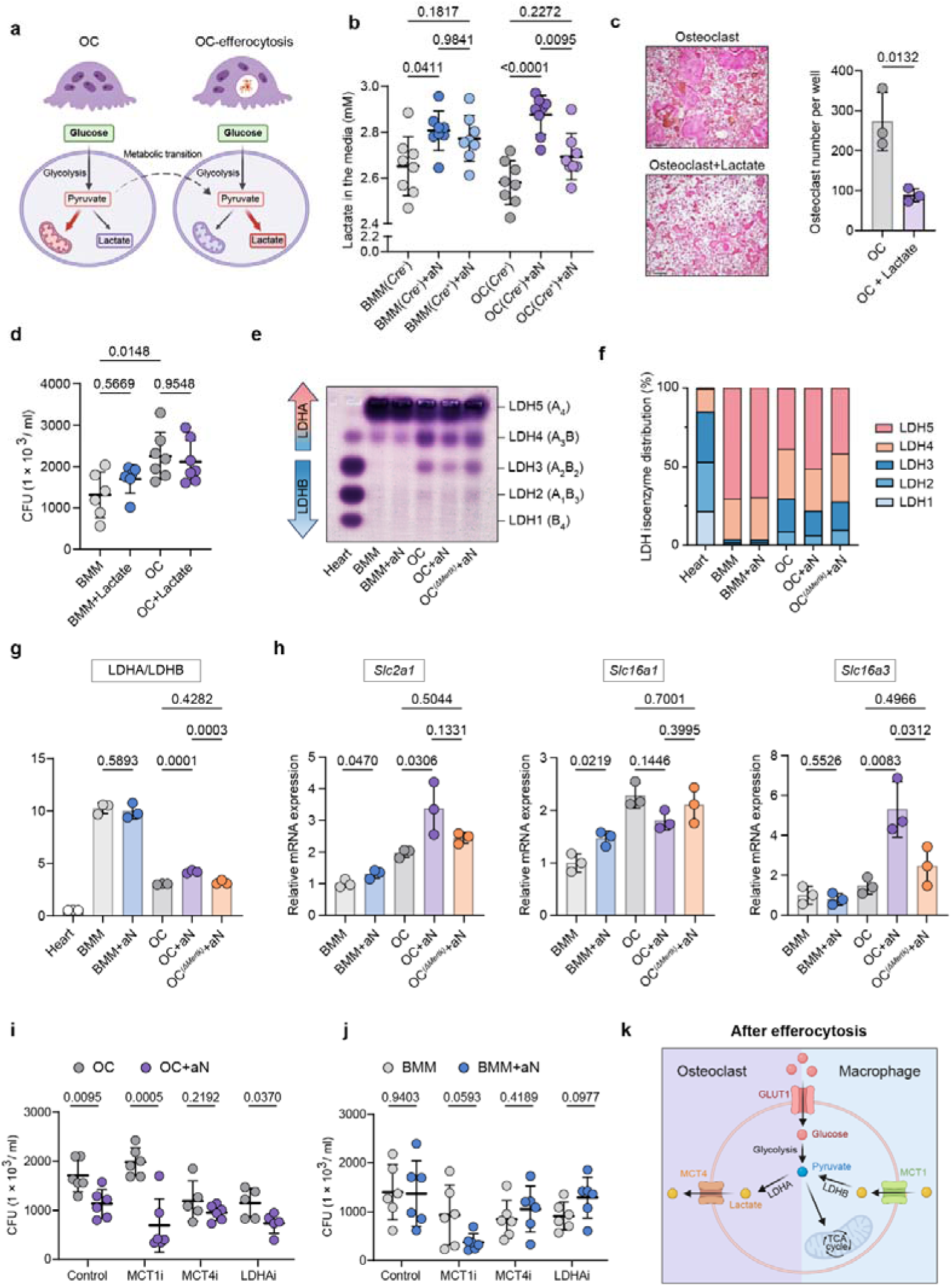
Lactate metabolic profiles and pharmacological responses in efferocytic osteoclasts and macrophages. **a**, Schematic illustrating osteoclast metabolic rewiring toward increased glycolytic activity after efferocytosis. **b**, Lactate concentration in the culture media of BMMs and osteoclasts derived from OC(ΔMertk) mice and control littermates before and after efferocytosis. **c,d**, Functional assessment of sodium lactate supplementation. Osteoclast differentiation was evaluated by TRAP staining (**c**). Scale bars, 100 µm. Antibacterial activity was assessed by CFU quantification after *Staphylococcus aureus* challenge (**d**). **e–g**, LDH isoenzyme profiling of BMMs and osteoclasts before and after efferocytosis. Representative agarose gel electrophoresis images showing LDH isoenzyme patterns detected by in-gel NAD⁺/NBT-PMS activity staining (**e**), densitometric quantification of individual isoenzymes (**f**), and activity-derived estimates of LDHA/LDHB ratios (**g**). **h**, RT–qPCR analysis of *Slc2a1*, *Slc16a1*, and *Slc16a3* expression before and after apoptotic neutrophil efferocytosis. Expression levels were normalized to *Actb*. **i,j**, Effects of pharmacological inhibition of MCT1, MCT4, and LDHA on viable bacterial counts in the presence or absence of apoptotic neutrophils in osteoclasts (**i**) and macrophages (**j**), assessed using CFU quantification after *Staphylococcus aureus* challenge. **k**, Proposed model of distinct lactate-handling programs in macrophages and osteoclasts after efferocytosis. The depicted transport directions and metabolic fates represent mechanistic hypotheses informed by the expression and isoenzyme profiles. Data are presented as mean ± s.d. Statistical significance was assessed using two-tailed unpaired *t*-tests (**c,i,j**) and one-way ANOVA followed by Tukey’s multiple comparisons test (**b,d,g,h**). n = 8 samples per group (b); n = 3 (**c,g,h**); n = 6–7 per group (**d**); n = 5–6 per group (**i**); n = 6 per group (**j**). Each point represents one independent biological sample.

Apoptotic neutrophil uptake increased lactate concentrations in the culture media of both macrophages and osteoclasts (Fig. 5b). However, exogenous lactate supplementation (10 mM) only partially recapitulated the effects of efferocytosis by reducing multinucleated osteoclast formation (Fig. 5c), without a significant reduction in viable bacterial counts or a coordinated induction of immune effector genes (Fig. 5d and Fig. S5a). Thus, extracellular lactate alone did not reproduce the antibacterial response associated with osteoclast efferocytosis.

We next compared the lactate metabolic profiles of osteoclasts and macrophages following apoptotic cell uptake. Lactate dehydrogenase (LDH) catalyzes the reversible conversion between pyruvate and lactate, with LDHA generally favoring lactate production and LDHB favoring lactate oxidation to pyruvate^38–42^. LDH exists as five tetrameric isoenzymes (LDH1–LDH5) composed of different combinations of LDHA and LDHB subunits. LDH isoenzyme analysis revealed different isoenzyme distributions in macrophages and osteoclasts: macrophages predominantly displayed LDH5, with lower levels of LDH4, whereas osteoclasts exhibited a broader isoenzyme spectrum with a greater contribution of LDH2–LDH4 activity (Fig. 5e, f), indicating a greater relative contribution of LDHB-containing isoenzymes^43,44^. Apoptotic neutrophil efferocytosis further remodeled the LDH isoenzyme profile of osteoclasts, resulting in an increased activity-derived estimate of the LDHA/LDHB ratio, a shift that was attenuated in Mertk-deficient osteoclast cultures (Fig. 5e–g). Consistent with these changes, apoptotic neutrophil uptake increased *Slc2a1* (encoding GLUT1) expression in both macrophages and osteoclasts, alongside distinct changes in lactate transporter expression. Efferocytic osteoclasts upregulated *Slc16a3* (encoding MCT4), whereas macrophages upregulated *Slc16a1* (encoding MCT1). In osteoclasts, qPCR showed a trend toward higher *Ldha* expression. These profiles point to different modes of lactate handling (Fig. 5h and Fig. S5b)^45–47^.

Functional interrogation revealed that lactate handling shapes the antibacterial consequences of efferocytosis. In osteoclast cultures, MCT1 blockade potentiated the antibacterial advantage associated with apoptotic neutrophil uptake, whereas MCT4 inhibition diminished this advantage. The aN-associated reduction in CFU persisted under LDHA inhibition (Figure 5i). In macrophages, apoptotic neutrophil uptake alone did not elicit a comparable antibacterial response, whereas MCT1 inhibition promoted bacterial clearance in aN-treated cultures; inhibition of MCT4 or LDHA produced more limited effects (Figure 5j). These findings connect distinct lactate metabolic programs to the functional divergence of osteoclast and macrophage efferocytosis. Whereas efferocytosis-derived lactate has been linked to immunoregulatory and pro-resolving responses in macrophages^9,26,36,37^, efferocytic osteoclasts exhibited increased *Slc16a3* expression and altered LDH isoenzyme activity alongside enhanced antibacterial defense. Together, these results support a model in which osteoclasts couple increased lactate production and export to antibacterial defense, while MCT1-mediated lactate handling restrains antibacterial activity (Figure 5k).

### MERTK in the osteoclast lineage limits bacterial burden and infection-associated bone loss

To determine whether osteoclast efferocytosis contributes to host defense during bone infection, we challenged *Ctsk*-Cre; *Mertk*^flox/flox^ mice and Cre^−^ littermate controls with *Staphylococcus aureus*. Compared with littermate controls, OC(ΔMertk) mice exhibited lower body weights during the 14-day infection period (Fig. 6a) and more severe cortical and trabecular bone destruction (Fig. 6b and Fig. S6a, b). OC(ΔMertk) mice also showed higher bacterial burden in femoral bone marrow samples (Fig. 6c).

**Fig. 6.**
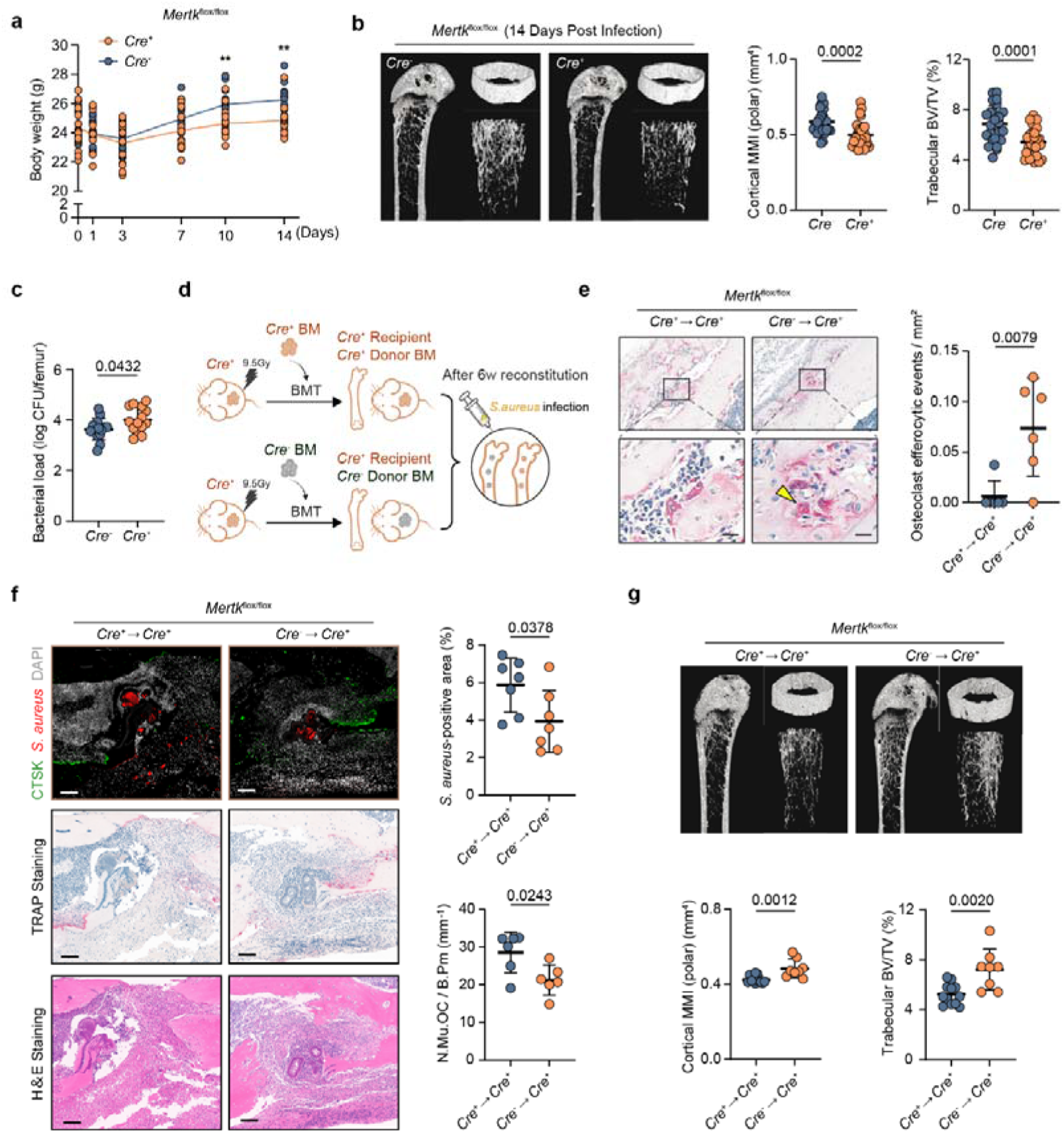
MERTK in the osteoclast lineage supports antibacterial defense and limits bone destruction. **a–c**, Phenotypic analysis of OC(ΔMertk) and littermate controls following *Staphylococcus aureus* bone infection. Body weight over 14 days (a), femoral micro-CT analysis at 14 dpi (b), and femoral bone marrow bacterial burden at 14 dpi (c). BV/TV, bone volume fraction; MMI (polar), mean polar moment of inertia. **d**, Schematic illustrating the bone marrow transplantation strategy. OC(ΔMertk) recipients were reconstituted with Cre^+^ or Cre^−^ bone marrow to generate Cre^+^ BM→Cre^+^ and Cre^−^ BM→Cre^+^ chimeric mice, respectively. **e**, TRAP staining and quantification of osteoclast-associated efferocytosis in femoral sections after BMT and *S. aureus* infection (14 dpi). Arrows indicate osteoclasts undergoing efferocytosis. Scale bars, 20 µm. **f**, Histological analysis of bone tissues from BMT-reconstituted mice at 14 dpi, with quantification of the *S. aureus*-positive areas (%) and osteoclast abundance. Scale bars, 100 µm. **g**, Micro-CT analysis of femurs from BMT-reconstituted mice at 14 dpi, with quantification of trabecular BV/TV and cortical MMI (polar). Data are presented as mean ± s.d. Statistical significance was assessed using two-tailed unpaired *t*-tests (**a,b,c,e,f,g**). n = 14 mice per genotype (**a,c**); n = 25 Cre− and 30 Cre+ mice (**b**); n = 6 mice per BMT group (**e**); n = 7 per group for S. aureus-positive area and n = 6 for osteoclast abundance (**f**); n = 12 Cre+ BM→Cre+ and 8 Cre− BM→Cre+ mice (**g**). Each point represents one mouse; measurements in a were repeated in the same animals.

Because *Ctsk*-Cre activity can also occur in non-osteoclast populations, including mesenchymal cells^48,49^, and MERTK signaling in these cells could potentially affect skeletal homeostasis, we next performed bone marrow transplantation (BMT) in OC(ΔMertk) recipients. The recipients were reconstituted with either Cre⁺ or Cre⁻ donor

BM, generating Cre⁺ BM→Cre⁺ and Cre⁻ BM→Cre⁺ chimeric mice, respectively (Fig. 6d). This strategy replaced the hematopoietic compartment while retaining recipient-derived mesenchymal cells^50^. Reconstitution with Cre⁻ BM restored MERTK expression in osteoclasts differentiated from recipient bone marrow (Fig. S6c) and increased osteoclast-associated efferocytosis in infected femoral sections (Fig. 6e).

Following *S. aureus* infection, reconstitution with Cre bone marrow reduced the *S. aureus*-positive area and osteoclast abundance at the infection site (Fig. 6f) and was accompanied by a reduction in the p-MLKL-positive area (Fig. S6d). Micro-CT analysis also showed less infection-associated bone destruction, with better preservation of trabecular and cortical bone architecture (Fig. 6g and Fig. S6e, f). Together with the preserved MERTK expression and efferocytosis in BMMs, these findings support a protective role for MERTK in the osteoclast lineage during bone infection.

## Discussion

Our study identifies an immune role for osteoclast efferocytosis during bone infection. Osteoclasts took up apoptotic neutrophils in infected bone, and exposure during differentiation reduced multinucleation and resorption while enhancing host-defense programs and antibacterial activity *in vitro*. *In vivo*, *Ctsk*-Cre-mediated *Mertk* deficiency increased bacterial burden and bone damage, whereas reconstitution with MERTK-competent bone marrow restored osteoclast efferocytosis and reduced infection-associated bone loss. These findings extend previous observations of apoptotic-cell uptake by osteoclasts^13,51^. During bone infection, immune pathways that control bacteria can also promote tissue damage. IL-1R/MyD88 signaling is required for local bacterial control but also promotes osteoclast accumulation and bone loss during infection^52^. Conversely, blocking RANKL-dependent osteoclast formation can prevent bone loss without lowering bacterial burden^53^. Here, apoptotic-neutrophil exposure reduced resorption while enhancing antibacterial activity in osteoclasts. Because bone contains protected bacterial niches and cannot be rapidly rebuilt^54^, this change in osteoclast function may be especially important at infected bone surfaces.

The outcome of efferocytosis is shaped by both apoptotic cargo and recipient-cell identity. In our experiments, the same apoptotic cargo elicited different responses in macrophages and osteoclasts. In macrophages, apoptotic-cell sensing can cooperate with reparative pathways, such as IL-4 or IL-13 signaling, to promote tissue remodeling^7,22^. Consistent with this context dependence, apoptotic neutrophil uptake increased CD206 expression in macrophages without a corresponding reduction in viable bacterial counts, whereas in osteoclasts it induced iNOS, host-defense programs, and antibacterial activity. These findings highlight the importance of recipient-cell identity and state in determining the outcome of efferocytosis. Although traditionally viewed as terminal effectors of bone resorption, osteoclasts also exhibit broader immune functions, including antigen presentation and regulation of T-cell responses^55,56^. In the setting of infection, conventional RANKL-differentiated osteoclasts can harbor intracellular *S. aureus* but display weaker bactericidal activity than their macrophage precursors^57^. Apoptotic-neutrophil exposure enhanced this antibacterial capacity. The distinct responses of macrophages and osteoclasts suggest that they could play complementary roles in infected bone, with osteoclasts linking local neutrophil turnover to host defense at bone surfaces.

Our metabolic findings link efferocytosis-associated metabolic remodeling to antibacterial function. Both macrophages and osteoclasts showed glycolytic responses to apoptotic-neutrophil uptake, yet a reduction in viable bacterial counts was observed only in osteoclast cultures. In osteoclasts, increased GLUT1 and HIF-1α expression, reduced oxygen consumption, higher extracellular lactate concentrations, and induction of *Slc16a3* together characterized the metabolic response. Deletion of *Mertk* blunted this metabolic shift, linking MERTK-mediated efferocytosis to metabolic remodeling. The induction of iNOS suggests a potential connection between immune signaling and metabolism, as nitric oxide can regulate mitochondrial metabolic pathways in inflammatory macrophages^58^. A similar interaction could help coordinate the immune and metabolic responses of efferocytic osteoclasts. Among the metabolic changes identified, lactate handling distinguished the responses of osteoclasts and macrophages. Exogenous lactate reduced osteoclast formation but did not reproduce the antibacterial phenotype.

MCT4-mediated lactate export may instead help sustain glycolytic activity; in macrophages, MCT4 loss impairs glycolytic remodeling and inflammatory output^59^. In our study, MCT4 and LDHA inhibition attenuated the antibacterial advantage associated with efferocytosis in osteoclast cultures, whereas MCT1 inhibition enhanced it. In macrophages, MCT1 inhibition lowered bacterial counts. The transporter and isoenzyme profiles, together with these functional responses, support a role for distinct lactate metabolic programs in shaping the antibacterial consequences of efferocytosis.

This study identifies efferocytosis as a mechanism that reshapes osteoclast function during infection. Signals derived from dying neutrophils coordinate reduced bone resorption with enhanced antibacterial activity, linking local neutrophil turnover to tissue preservation and host defense. Efferocytosis therefore represents an infection-adaptive component of the osteoclast immune repertoire. More broadly, our findings extend the functional scope of efferocytosis beyond cellular disposal, revealing apoptotic-cell uptake as a context-dependent signal that shapes the metabolic and immune responses of specialized phagocytes.

## Data availability

The RNA-sequencing data generated in this study are being deposited in a public repository and will be made publicly available upon publication. Accession numbers and repository links will be provided once deposition is complete. Source data underlying the figures will be made available with the published article. Accession numbers for the publicly available datasets reanalyzed in this study are provided in the Methods and figure legends.

## Acknowledgments

This work was supported by the National Natural Science Foundation of China (82572785), the NSFC Key Projects of the Regional Innovation and Development Joint Fund (U23A20413), and the Southwest Hospital Boqing Innovation Fund (2024BQCXJJ-9).

## Author Contributions

Conceptualization: C.D., F.L., Q.Y.;

Methodology: Q.Y., Y.Q., J.Z.;

Investigation: Q.Y., Y.Q., J.Z., Y.J., Y.T., Y.Z.;

Formal Analysis: J.Z., Q.Y., Y.Q., Q.D., Z.H.;

Resources: Y.Q., J.Z., F.L., Y.J., Q.D., J.S.;

Writing -Original Draft: Q.Y., J.Z.;

Writing -Review & Editing: Q.Y., N.L., F.L., C.D.;

Visualization: J.Z., Y.Q., C.D.;

Supervision: F.L., C.D.;

Project Administration: C.D.;

Funding Acquisition: F.L., J.Z., C.D.

## Declaration of Interests

The authors declare no competing interests.

## Materials and methods

### Mice

All animal procedures were conducted in accordance with institutional ethical guidelines and were approved by the Institutional Animal Care and Use Committee of Army Medical University. All mouse strains used in this study were maintained on the C57BL/6 background. Male mice were used throughout the study. Non-transplanted mice were 8–10 weeks old at the start of experiments; the timing of experiments in bone marrow transplant recipients is described below. C57BL/6J mice were obtained from the Southwest Hospital Animal Center (Chongqing, China), whereas Ly6G-tdTomato mice and *Ctsk*-Cre; *Mertk*^flox/flox^ mice were obtained from Cyagen Biotechnology Co., Ltd. (Suzhou, China). Mice were housed under specific pathogen-free conditions at a controlled temperature on a 12-h light/12-h dark cycle, with ad libitum access to food and water.

### Cell culture

Cell cultures were maintained under aseptic conditions, with *Staphylococcus aureus* introduced only for the indicated bacterial challenge assays. Mouse neutrophils were freshly isolated, and primary macrophages and osteoclasts were generated as described below. Unless otherwise indicated, cells were maintained at 37°C in an atmosphere containing 5% CO₂. The murine macrophage-like cell line RAW264.7 (ATCC TIB-71; RRID: CVCL_0493) was cultured according to the manufacturer’s protocols.

### Neutrophil isolation and induction of apoptosis

Bone marrow neutrophils were isolated from mice by negative selection using the EasySep™ Mouse Neutrophil Enrichment Kit (STEMCELL Technologies) according to the manufacturer’s instructions. Purity was assessed by flow cytometry, with neutrophils defined as Ly6G⁺CD11b⁺ cells. Cell preparations with a purity greater than 90% were used. To induce apoptosis, freshly isolated neutrophils were cultured in RPMI-1640 medium supplemented with 2–5% fetal bovine serum for 24 h at 37°C to allow spontaneous apoptosis.

### *In vitro* BMM generation and osteoclast differentiation

Bone marrow cells were flushed from the femurs and tibias of mice, subjected to red blood cell lysis, and passed through a 70-μm cell strainer. Cells were cultured in αMEM supplemented with 10% FBS and 1% penicillin–streptomycin. After 24 h, nonadherent cells were collected and cultured with M-CSF (25 ng/mL; R&D Systems) for 2 d to generate bone marrow-derived macrophages (BMMs). To induce osteoclast differentiation, BMMs were subsequently cultured with M-CSF (25 ng/mL) and RANKL (50 ng/mL; R&D Systems) for an additional 4–5 d until multinucleated osteoclasts were observed. Where indicated, cells were treated with sodium lactate (10 mM; Aladdin), TNF-α (10 ng/mL; TargetMol), IL-4 (10 ng/mL; TargetMol), the MCT1 inhibitor AZD3965 (100 nM; TargetMol), the MCT4 inhibitor VB124 (10 μM; TargetMol), or the LDHA inhibitor FX11 (10 μM; TargetMol).

Unless otherwise specified, for efferocytosis experiments, apoptotic neutrophils were added to BMM cultures or to osteoclast cultures after 3 d of RANKL stimulation at a ratio of 5:1 (apoptotic neutrophils to phagocytes). Control cultures received an equal volume of PBS. Cells were collected for subsequent analyses when multinucleated osteoclasts became apparent in the corresponding control cultures.

### Murine model of acute osteomyelitis

A murine model of acute osteomyelitis was established as previously described^14,15^. Briefly, non-transplanted mice aged 8–10 weeks, or bone marrow transplant recipients after 6 weeks of hematopoietic reconstitution, were anesthetized with inhaled isoflurane, and a unicortical bone defect was created in the proximal femur using a 29-gauge needle to generate a minimally invasive microchannel.

Subsequently, 2 μL of PBS containing 1 × 10 CFU of *Staphylococcus aureus* ATCC 43300 was injected into the femoral medullary canal through the defect using a 2.5 μL Hamilton microsyringe equipped with a 33-gauge needle.

Except for longitudinal studies, for which assessment time points are specified in the corresponding sections, micro-CT, bacterial burden, and histological analyses were performed on day 14 post-infection. Because bacterial injection through the proximal femoral defect caused localized cortical bone damage at the injection site, micro-CT analysis of cortical and trabecular bone was restricted to distal femoral regions of interest remote from the surgical site.

### Flow cytometric analysis

Bone marrow cells were harvested from the femurs of mice, filtered through 70-μm cell strainers, and treated with ACK lysis buffer for 5 min at room temperature to remove red blood cells. Cells were resuspended in Annexin V binding buffer at a density of 1 × 10 cells/mL. Aliquots of 100 μL of cell suspension were incubated with anti-Ly6G-APC antibody (BioLegend) for 30 min at 4°C in the dark, followed by staining with Annexin V-FITC and propidium iodide (PI) (BioLegend) for 15 min at room temperature in the dark. Samples were analyzed by flow cytometry, and data were processed using FlowJo software.

For efferocytosis assays, phagocytes and apoptotic neutrophils were labeled with DiI and DiO (Sangon Biotech; E607331 and E607332, respectively) to distinguish the two cell populations. After labeling, cells were washed thoroughly and co-cultured for 1 h to allow efferocytosis. Following co-culture, cells were washed to remove non-engulfed apoptotic neutrophils and collected for flow cytometric analysis to quantify the uptake of apoptotic neutrophils by phagocytes. Efferocytosis was quantified as the percentage of DiI⁺ phagocytes that acquired the DiO signal from apoptotic neutrophils. Because the percentage of apoptotic cell uptake varied among independent experiments, a normalized phagocytosis index was calculated as uptake in the experimental group (%) divided by uptake in the control group (%).

### Immunofluorescence staining

Femurs were collected, fixed in 4% paraformaldehyde, decalcified, and embedded in paraffin. Serial paraffin sections were prepared, deparaffinized, rehydrated, subjected to antigen retrieval, and blocked. Sections were incubated overnight at 4°C with primary antibodies against Ly6G (Servicebio; GB11229), CTSK (Santa Cruz Biotechnology; sc-48353), MERTK (Proteintech; 27900-1-AP), AXL (MedChemExpress; HY-P81198), phospho-MLKL (Ser345) (Abcam; ab196436), and *Staphylococcus aureus* (Abcam; ab20920), followed by incubation with the corresponding fluorophore-conjugated secondary antibodies. TUNEL staining (Servicebio; G1504) was performed according to the manufacturer’s instructions to detect apoptotic cells. For cellular immunofluorescence staining, cells cultured on glass coverslips were fixed with 4% paraformaldehyde, permeabilized, and blocked. Cells were incubated overnight at 4°C with primary antibodies against Vinculin (Abcam; ab91459), iNOS (Santa Cruz Biotechnology; sc-7271), and CD206 (ABclonal; A21014). After washing, cells were incubated with the corresponding fluorophore-conjugated secondary antibodies. F-actin was visualized using fluorescently conjugated phalloidin staining (Abcam; ab176753). Nuclei were counterstained with DAPI. Fluorescence signals were quantified using ImageJ.

### Western blot analysis

Lysates were separated by 10% SDS-PAGE and transferred onto PVDF membranes with a 0.45-μm pore size. Membranes were blocked with 5% nonfat milk in TBST (TBS containing 0.1% Tween-20) for 1 h at room temperature and then incubated overnight at 4°C with the indicated primary antibodies against GAS6 (MedChemExpress; HY-P85881), MERTK (Proteintech; 27900-1-AP), AXL (MedChemExpress; HY-P81198), HIF-1α (Proteintech; 82989-4-RR), GLUT1 (Proteintech; 21829-1-AP), and OXPHOS Cocktail (Proteintech; PK30006). After three washes with TBST, membranes were incubated with HRP-conjugated secondary antibodies (Affinity Biosciences) for 1 h at room temperature. Immunoreactive bands were visualized using chemiluminescence reagents (ECL; Thermo Fisher Scientific) and imaged using a ChemiDoc Imaging System (Bio-Rad). After stripping, membranes were reprobed with an HRP-conjugated recombinant anti-β-actin antibody (ServiceBio; ZB15001-HRP-100). Band intensities were quantified using ImageJ and normalized to β-actin.

### TRAP staining and histological analysis

For *in vitro* osteoclast differentiation assays, cells were fixed with 4% paraformaldehyde for 10 min and stained using a tartrate-resistant acid phosphatase (TRAP) staining kit (GENMED) according to the manufacturer’s instructions. TRAP⁺ multinucleated cells containing at least three nuclei were identified as osteoclasts and quantified by light microscopy.

For histological analysis, decalcified femoral sections were subjected to hematoxylin and eosin (H&E) staining (Servicebio; G1005), Safranin O/Fast Green staining (Servicebio; G1053), and TRAP staining (GENMED) according to the manufacturers’ instructions. TRAP⁺ multinucleated osteoclasts along the trabecular bone perimeter in the distal femur were quantified and expressed as the number of multinucleated osteoclasts per millimeter of bone perimeter (N.Mu.OC/B.Pm). Histological images were acquired using light microscopy and analyzed using ImageJ.

### Bone resorption assay and scanning electron microscopy

Bovine cortical bone slices (Boneslices.com, Jelling, Denmark) were placed in 96-well plates. Osteoclast precursors were seeded onto the bone slices and cultured as described above, with parallel wells without bone slices included as controls to monitor osteoclast formation. Culture medium containing M-CSF and RANKL was replaced every 2 d. After multinucleated osteoclasts were observed in the parallel control wells, cells cultured on bone slices were maintained for an additional 7 d to allow bone resorption. Bone slices were then removed from the wells and washed thoroughly to remove residual cells. For scanning electron microscopy, bone slices were fixed, dehydrated through a graded ethanol series, air-dried, mounted on specimen stubs, and sputter-coated with gold. Resorption pits were imaged using a scanning electron microscope.

### Bulk RNA sequencing and single-cell transcriptomic analysis

For bulk RNA-seq analysis, BMMs and osteoclasts were generated *in vitro* as described above and incubated with apoptotic neutrophils. After a 2-h efferocytosis pulse, non-engulfed apoptotic neutrophils were removed, and the cultures were further incubated for 12 h before sample collection. Three biological replicates were prepared for each group. Total RNA was extracted using TRIzol reagent (Thermo Fisher Scientific) according to the manufacturer’s instructions. Poly(A)-enriched, strand-specific mRNA libraries were prepared through RNA fragmentation, reverse transcription, second-strand cDNA synthesis, adapter ligation, and PCR amplification. Libraries that passed quality control were sequenced on an Illumina NovaSeq X Plus platform to generate 150-bp paired-end reads. Differential expression analysis was performed using DESeq2. Genes with an adjusted *P* value < 0.05 and |log₂(fold change)| ≥ 1 were considered differentially expressed.

To evaluate the transcriptional similarity of macrophages, osteoclasts, and efferocytic osteoclasts to established macrophage activation states, the publicly available scRNA-seq dataset GSE117176 was analyzed using Seurat in R.

State-specific gene signatures for macrophages, osteoclasts, and efferocytic osteoclasts were defined as the 500 genes with the highest state-specificity scores for each group based on the bulk RNA-seq profiles. Module scores for these signatures were calculated using the AddModuleScore function and visualized on a t-SNE embedding. Normalized module scores were subsequently used to calculate the relative transcriptional affinity of individual cells for the three reference states.

Transcriptional similarities to macrophage activation states were further characterized using MacSpectrum^29^, which integrates the macrophage polarization index (MPI) and activation-induced macrophage differentiation index (AMDI) to quantify macrophage polarization and activation-associated differentiation based on transcriptomic profiles.

### Reverse transcription–quantitative PCR

For RT-qPCR analysis, total RNA was extracted using TRIzol reagent (Invitrogen) according to the manufacturer’s instructions. RNA concentration and purity were assessed using a NanoDrop 2000 spectrophotometer (Thermo Fisher Scientific). One microgram of total RNA was reverse-transcribed into cDNA using the PrimeScript RT Reagent Kit with gDNA Eraser (Takara). Quantitative PCR was performed using TB Green Premix Ex Taq (Takara) on a CFX96 Touch Real-Time PCR Detection System (Bio-Rad) according to the manufacturer’s protocol. *Actb* was used as the internal reference gene. Primer sequences are provided in Table S1.

### siRNA-mediated gene silencing

For gene-silencing experiments, BMMs were transfected with siRNAs targeting *Axl* (si-*Axl*) or *Mertk* (si-*Mertk*), or with non-targeting control siRNA (si-NC) (Hanbio). siRNAs were transfected at a final concentration of 20 nM using RNAFit transfection reagent (Hanbio) according to the manufacturer’s instructions. After transfection, cells were cultured for 72 h before subsequent experiments. Knockdown efficiency was confirmed by RT-qPCR. The siRNA sequences are provided in Table S2.

### Tissue and cell-based CFU enumeration

For tissue CFU enumeration, femurs were aseptically collected and cleared of surrounding soft tissues. After removal of both epiphyses, the bone marrow was flushed with a defined volume of sterile PBS into prechilled tubes. Bone marrow suspensions were serially diluted in sterile PBS and plated on tryptic soy agar (TSA) plates. After incubation at 37 °C for 24 h, bacterial colonies were counted, and bacterial burden was expressed as CFU per femur.

For cell-based CFU enumeration, BMMs and osteoclasts were subjected to the indicated treatments before challenge with *Staphylococcus aureus* at a multiplicity of infection (MOI) of 10. After incubation for 12 h, viable bacteria were recovered from the cell cultures, serially diluted in sterile PBS, and plated on tryptic soy agar (TSA) plates. After incubation at 37 °C for 24 h, bacterial colonies were counted, and viable bacterial counts were expressed as CFU/mL.

### Seahorse metabolic analysis

Cellular metabolic profiles were analyzed using a Seahorse XF96 Analyzer (Agilent Technologies). BMMs were seeded in XF96 cell culture microplates. For osteoclast assays, BMMs were seeded and differentiated into osteoclasts by stimulation with M-CSF and RANKL. For efferocytosis assays, apoptotic neutrophils were added at a ratio of 5:1 (apoptotic neutrophils to phagocytes). After a 2-h efferocytosis pulse, non-engulfed apoptotic neutrophils were removed, and the cells were incubated for an additional 12 h before metabolic analysis.

For the mitochondrial stress test, oligomycin (2 μM), FCCP (1 μM), and rotenone/antimycin A (0.5 μM each) were sequentially injected, and the oxygen consumption rate (OCR) was measured. For the glycolysis stress test, glucose (10 mM), oligomycin (2 μM), and 2-deoxy-d-glucose (2-DG; 50 mM) were sequentially injected, and the extracellular acidification rate (ECAR) was measured. Data were acquired and analyzed using Seahorse Wave software, and the corresponding metabolic parameters were calculated according to the manufacturer’s instructions.

### Micro-computed tomography analysis

Bone architecture in *Staphylococcus aureus*–infected femurs was assessed by high-resolution microcomputed tomography using a SkyScan 1272 system (Bruker, Kontich, Belgium). Femurs were fixed in 4% paraformaldehyde at 4 °C for 72 h, thoroughly rinsed with PBS, and scanned at 80 kV and 165 μA with an isotropic voxel size of 7 μm. A 4-mm-long region of interest (ROI) encompassing the distal femoral metaphysis was selected using identical anatomical landmarks for all samples. Raw projection images were reconstructed using NRecon software (Bruker), and morphometric parameters were computed with CT Analyzer (CTAn, v1.18.8; Bruker). Three-dimensional anatomical renderings were generated using CTVol software (Bruker), and osteolytic lesions were visually assessed in reconstructed coronal and sagittal planes.

### Lactate measurement and LDH isoenzyme analysis

Lactate concentrations in cell culture supernatants were measured using an L-Lactate Assay Kit with WST-8 (Beyotime, S0208S) according to the manufacturer’s instructions. LDH isoenzyme activity was analyzed by non-denaturing agarose gel electrophoresis followed by in-gel enzymatic activity staining. Briefly, cell lysates were separated on agarose gels, and LDH isoenzymes were visualized using a staining solution containing lactate, NAD⁺, nitroblue tetrazolium (NBT), and phenazine methosulfate (PMS). LDH1–LDH5 bands were identified according to their electrophoretic mobility, and band intensities were quantified using ImageJ software. The relative activity of each LDH isoenzyme was calculated as its band intensity divided by the total intensity of all five LDH isoenzyme bands. An activity-derived estimate of the LDHA-to-LDHB subunit ratio was calculated from the weighted distribution of LDH1–LDH5 isoenzymes.

### Bone marrow transplantation

Recipient mice received myeloablative total-body irradiation at a total dose of 9.5 Gy using a ^60^Co γ-irradiator. Bone marrow cells were aseptically isolated from the femurs and tibias of donor mice and passed through a 70-μm cell strainer. A total of 5 × 10 donor bone marrow cells were administered intravenously to each recipient through the tail vein. The recipients were allowed to recover for 6 weeks to permit hematopoietic reconstitution before subsequent experiments. Restoration of MERTK expression in the osteoclast lineage was assessed by measuring MERTK protein abundance in osteoclasts differentiated from recipient bone marrow cells. Recipients were 8–10 weeks old at transplantation and 14–16 weeks old at infection after the 6-week recovery period; tissues collected at 14 dpi were therefore obtained at 16–18 weeks of age.

### Statistics

Data are presented as mean ± s.d. Statistical analyses were performed using GraphPad Prism. Comparisons between two groups were conducted using two-tailed unpaired Student’s *t*-tests. Comparisons among three or more groups were performed using one-way analysis of variance (ANOVA) followed by Tukey’s multiple-comparisons test.

## Supplemental material

Figs. S1–S6 provide additional characterization of osteoclast efferocytosis, metabolism, and infection-associated phenotypes. Tables S1 and S2 contain primer and siRNA information, respectively, and are supplied as separate Excel files.

## Notes

### Competing Interest Statement

The authors have declared no competing interest.

## References

1. P. Mehrotra, K. S. Ravichandran, Drugging the efferocytosis process: concepts and opportunities. Nat. Rev. Drug Discov. 21, 601–620 (2022).

2. A. C. Doran, A. Yurdagul, Jr., I. Tabas, Efferocytosis in health and disease. Nat. Rev. Immunol. 20, 254–267 (2020).

3. M. R. Elliott, K. S. Ravichandran, The Dynamics of Apoptotic Cell Clearance. Dev. Cell 38, 147–160 (2016).

4. E. Boada-Romero, J. Martinez, B. L. Heckmann, D. R. Green, The clearance of dead cells by efferocytosis. Nat. Rev. Mol. Cell Biol. 21, 398–414 (2020).

5. S. Morioka, C. Maueroder, K. S. Ravichandran, Living on the Edge: Efferocytosis at the Interface of Homeostasis and Pathology. Immunity 50, 1149–1162 (2019).

6. L. Mukundan, J. I. Odegaard, C. R. Morel, J. E. Heredia, J. W. Mwangi, R. R. Ricardo-Gonzalez, Y. P. S. Goh, A. R. Eagle, S. E. Dunn, J. U. H. Awakuni, K. D. Nguyen, L. Steinman, S. A. Michie, A. Chawla, PPAR-delta senses and orchestrates clearance of apoptotic cells to promote tolerance. Nat. Med. 15, 1266–1272 (2009).

7. L. Bosurgi, Y. G. Cao, M. Cabeza-Cabrerizo, A. Tucci, L. D. Hughes, Y. Kong, J. S. Weinstein, P. Licona-Limon, E. T. Schmid, F. Pelorosso, N. Gagliani, J. E. Craft, R. A. Flavell, S. Ghosh, C. V. Rothlin, Macrophage function in tissue repair and remodeling requires IL-4 or IL-13 with apoptotic cells. Science 356, 1072–1076 (2017).

8. S. Zhang, S. Weinberg, M. DeBerge, A. Gainullina, M. Schipma, J. M. Kinchen, I. Ben-Sahra, D. R. Gius, L. Yvan-Charvet, N. S. Chandel, P. T. Schumacker, E. B. Thorp, Efferocytosis Fuels Requirements of Fatty Acid Oxidation and the Electron Transport Chain to Polarize Macrophages for Tissue Repair. Cell Metab. 29, 443–456.e5 (2019).

9. S. Morioka, J. S. A. Perry, M. H. Raymond, C. B. Medina, Y. Zhu, L. Zhao, V. Serbulea, S. Onengut-Gumuscu, N. Leitinger, S. Kucenas, J. C. Rathmell, L. Makowski, K. S. Ravichandran, Efferocytosis induces a novel SLC program to promote glucose uptake and lactate release. Nature 563, 714–718 (2018).

10. C. E. Jacome-Galarza, G. I. Percin, J. T. Muller, E. Mass, T. Lazarov, J. Eitler, M. Rauner, V. K. Yadav, L. Crozet, M. Bohm, P. L. Loyher, G. Karsenty, C. Waskow, F. Geissmann, Developmental origin, functional maintenance and genetic rescue of osteoclasts. Nature 568, 541–545 (2019).

11. J. W. Lee, I. H. Lee, T. Iimura, S. W. Kong, Two macrophages, osteoclasts and microglia: from development to pleiotropy. Bone Res. 9, 11 (2021).

12. S. Lemma, M. Sboarina, P. E. Porporato, N. Zini, P. Sonveaux, G. Di Pompo, N. Baldini, S. Avnet, Energy metabolism in osteoclast formation and activity. Int. J. Biochem. Cell Biol. 79, 168–180 (2016).

13. U. Harre, H. Keppeler, N. Ipseiz, A. Derer, K. Poller, M. Aigner, G. Schett, M. Herrmann, K. Lauber, Moonlighting osteoclasts as undertakers of apoptotic cells. Autoimmunity 45, 612–619 (2012).

14. J. Zhang, Q. Yu, Y. Qu, J. Tan, Q. Dai, Y. Jia, J. Xu, Y. Yang, Y. Tang, Y. Zhang, S. Guo, Z. Zhang, Y. Zou, N. Limjunyawong, F. Luo, C. Dou, Mrgpra2(+) neutrophils integrate infection-derived signals to trigger NET-mediated antimicrobial defense in bone marrow. Cell Rep. 45, 117579 (2026).

15. Q. Yu, Y. Qu, J. Zhang, Z. Zhang, S. Guo, F. Luo, C. Dou, Neutrophil-derived apoptotic bodies function as decoy signals to preserve bone integrity during infectious osteomyelitis. Bone Res. 14, 90 (2026).

16. C. V. Rothlin, E. A. Carrera-Silva, L. Bosurgi, S. Ghosh, TAM receptor signaling in immune homeostasis. Annu. Rev. Immunol. 33, 355–391 (2015).

17. K. Nagata, K. Ohashi, T. Nakano, H. Arita, C. Zong, H. Hanafusa, K. Mizuno, Identification of the product of growth arrest-specific gene 6 as a common ligand for Axl, Sky, and Mer receptor tyrosine kinases. J. Biol. Chem. 271, 30022–30027 (1996).

18. Y. Morita, M. Saito, J. Rangel-Moreno, A. M. Franchini, J. R. Owen, J. C. Martinez, J. L. Daiss, K. L. de Mesy Bentley, S. L. Kates, E. M. Schwarz, G. Muthukrishnan, Systemic IL-27 administration prevents abscess formation and osteolysis via local neutrophil recruitment and activation. Bone Res. 10, 56 (2022).

19. T. Burstyn-Cohen, R. Fresia, TAM receptors in phagocytosis: Beyond the mere internalization of particles. Immunol. Rev. 319, 7–26 (2023).

20. M. S. Hansen, K. Madsen, M. Price, K. Søe, Y. Omata, M. M. Zaiss, C. M. Gorvin, M. Frost, A. Rauch, Transcriptional reprogramming during human osteoclast differentiation identifies regulators of osteoclast activity. Bone Res. 12, 5 (2024).

21. M. Tsukasaki, N. C. N. Huynh, K. Okamoto, R. Muro, A. Terashima, Y. Kurikawa, N. Komatsu, W. Pluemsakunthai, T. Nitta, T. Abe, H. Kiyonari, T. Okamura, M. Sakai, T. Matsukawa, M. Matsumoto, Y. Kobayashi, J. M. Penninger, H. Takayanagi, Stepwise cell fate decision pathways during osteoclastogenesis at single-cell resolution. Nat. Metab. 2, 1382–1390 (2020).

22. I. Liebold, A. Al Jawazneh, C. Casar, C. Lanzloth, S. Leyk, M. Hamley, M. N. Wong, D. Kylies, S. K. Gräfe, I. Edenhofer, I. Aranda-Pardos, M. Kriwet, H. Haas, J. Krause, A. Hadjilaou, A. B. Schromm, U. Richardt, P. Eggert, D. Tappe, S. A. Weidemann, S. Ghosh, C. F. Krebs, N. A-Gonzalez, A. Worthmann, A. W. Lohse, S. Huber, C. V. Rothlin, V. G. Puelles, T. Jacobs, N. Gagliani, L. Bosurgi, Apoptotic cell identity induces distinct functional responses to IL-4 in efferocytic macrophages. Science 384, eabo7027 (2024).

23. G. Lemke, How macrophages deal with death. Nat. Rev. Immunol. 19, 539–549 (2019).

24. M. C. Greenlee-Wacker, Clearance of apoptotic neutrophils and resolution of inflammation. Immunol. Rev. 273, 357–370 (2016).

25. S. Maschalidi, P. Mehrotra, B. N. Keçeli, H. K. L. De Cleene, K. Lecomte, R. Van der Cruyssen, P. Janssen, J. Pinney, G. van Loo, D. Elewaut, A. Massie, E. Hoste, K. S. Ravichandran, Targeting SLC7A11 improves efferocytosis by dendritic cells and wound healing in diabetes. Nature 606, 776–784 (2022).

26. D. Ngai, M. Schilperoort, I. Tabas, Efferocytosis-induced lactate enables the proliferation of pro-resolving macrophages to mediate tissue repair. Nat. Metab. 5, 2206–2219 (2023).

27. S. R. Sukka, P. B. Ampomah, L. N. F. Darville, D. Ngai, X. Wang, G. Kuriakose, Y. Xiao, J. Shi, J. M. Koomen, R. H. McCusker, I. Tabas, Efferocytosis drives a tryptophan metabolism pathway in macrophages to promote tissue resolution. Nat. Metab. 6, 1736–1755 (2024).

28. M. R. Elliott, F. B. Chekeni, P. C. Trampont, E. R. Lazarowski, A. Kadl, S. F. Walk, D. Park, R. I. Woodson, M. Ostankovich, P. Sharma, J. J. Lysiak, T. K. Harden, N. Leitinger, K. S. Ravichandran, Nucleotides released by apoptotic cells act as a find-me signal to promote phagocytic clearance. Nature 461, 282–286 (2009).

29. C. Li, A. Menoret, C. Farragher, Z. Ouyang, C. Bonin, P. Holvoet, A. T. Vella, B. Zhou, Single-cell transcriptomics–based MacSpectrum reveals macrophage activation signatures in diseases. JCI Insight 4, e126453 (2019).

30. P. Ettel, T. Weichhart, Not just sugar: metabolic control of neutrophil development and effector functions. J. Leukoc. Biol. 116, 487–510 (2024).

31. X. Chen, L. Liu, S. Kang, J. R. Gnanaprakasam, R. Wang, The lactate dehydrogenase (LDH) isoenzyme spectrum enables optimally controlling T cell glycolysis and differentiation. Sci. Adv. 9, eadd9554 (2023).

32. L. A. O’Neill, R. J. Kishton, J. Rathmell, A guide to immunometabolism for immunologists. Nat. Rev. Immunol. 16, 553–565 (2016).

33. S. E. Corcoran, L. A. O’Neill, HIF1alpha and metabolic reprogramming in inflammation. J. Clin. Invest. 126, 3699–3707 (2016).

34. G. M. Tannahill, A. M. Curtis, J. Adamik, E. M. Palsson-McDermott, A. F. McGettrick, G. Goel, C. Frezza, N. J. Bernard, B. Kelly, N. H. Foley, L. Zheng, A. Gardet, Z. Tong, S. S. Jany, S. C. Corr, M. Haneklaus, B. E. Caffrey, K. Pierce, S. Walmsley, F. C. Beasley, E. Cummins, V. Nizet, M. Whyte, C. T. Taylor, H. Lin, S. L. Masters, E. Gottlieb, V. P. Kelly, C. Clish, P. E. Auron, R. J. Xavier, L. A. J. O’Neill, Succinate is an inflammatory signal that induces IL-1beta through HIF-1alpha. Nature 496, 238–242 (2013).

35. A. J. Freemerman, A. R. Johnson, G. N. Sacks, J. J. Milner, E. L. Kirk, M. A. Troester, A. N. Macintyre, P. Goraksha-Hicks, J. C. Rathmell, L. Makowski, Metabolic reprogramming of macrophages: glucose transporter 1 (GLUT1)-mediated glucose metabolism drives a proinflammatory phenotype. J. Biol. Chem. 289, 7884–7896 (2014).

36. M. Schilperoort, D. Ngai, M. Katerelos, D. A. Power, I. Tabas, PFKFB2-mediated glycolysis promotes lactate-driven continual efferocytosis by macrophages. Nat. Metab. 5, 431–444 (2023).

37. R. Kannan, N. J. Carruthers, A. J. Koh, G. G. Kleer, K. Nakamura, S. C. J. Parker, L. K. McCauley, H. Roca, Efferocytosis-induced metabolic shift in bone macrophages drives lactate production and modulates inflammation and osteoclastogenesis. Front. Immunol. 16, 1650465 (2025).

38. D. M. Dawson, T. L. Goodfriend, N. O. Kaplan, Lactic Dehydrogenases: Functions of the Two Types Rates of Synthesis of the Two Major Forms Can Be Correlated with Metabolic Differentiation. Science 143, 929–933 (1964).

39. A. A. Khan, K. S. Allemailem, F. A. Alhumaydhi, S. J. T. Gowder, A. H. Rahmani, The Biochemical and Clinical Perspectives of Lactate Dehydrogenase: An Enzyme of Active Metabolism. Endocr. Metab. Immune Disord. Drug Targets 20, 855–868 (2020).

40. J. M. Ross, J. Öberg, S. Brené, G. Coppotelli, M. Terzioglu, K. Pernold, M. Goiny, R. Sitnikov, J. Kehr, A. Trifunovic, N. G. Larsson, B. J. Hoffer, L. Olson, High brain lactate is a hallmark of aging and caused by a shift in the lactate dehydrogenase A/B ratio. Proc. Natl. Acad. Sci. U.S.A. 107, 20087–20092 (2010).

41. K. Swiderek, P. Paneth, Differences and similarities in binding of pyruvate and L-lactate in the active site of M4 and H4 isoforms of human lactate dehydrogenase. Arch. Biochem. Biophys. 505, 33–41 (2011).

42. J. A. Read, V. J. Winter, C. M. Eszes, R. B. Sessions, R. L. Brady, Structural basis for altered activity of M-and H-isozyme forms of human lactate dehydrogenase. Proteins 43, 175–185 (2001).

43. H. Ahn, K. Lee, J. M. Kim, S. H. Kwon, S. H. Lee, S. Y. Lee, D. Jeong, Accelerated Lactate Dehydrogenase Activity Potentiates Osteoclastogenesis via NFATc1 Signaling. PLoS One 11, e0153886 (2016).

44. S. Lemma, G. Di Pompo, P. E. Porporato, M. Sboarina, S. Russell, R. J. Gillies, N. Baldini, P. Sonveaux, S. Avnet, MDA-MB-231 breast cancer cells fuel osteoclast metabolism and activity: A new rationale for the pathogenesis of osteolytic bone metastases. Biochim. Biophys. Acta Mol. Basis Dis. 1863, 3254–3264 (2017).

45. A. P. Halestrap, M. C. Wilson, The monocarboxylate transporter family--role and regulation. IUBMB Life 64, 109–119 (2012).

46. A. A. Cluntun, J. R. Visker, J. N. Velasco-Silva, M. J. Lang, L. Cedeno-Rosario, T. S. Shankar, R. Hamouche, J. Ling, J. E. Kim, A. G. Toshniwal, H. K. Low, C. N. Cunningham, J. Carrington, J. L. Catrow, Q. Pearce, M. Y. Jeong, A. J. Bott, A. J. Narbona-Perez, C. E. Stanley, Q. Li, D. R. Eberhardt, J. T. Morgan, T. Yadav, C. E. Wells, D. K. A. Ramadurai, W. I. Swiatek, D. Chaudhuri, J. D. Rothstein, D. M. Muoio, J. A. Paulo, S. P. Gygi, S. A. Baker, S. Navankasattusas, J. E. Cox, K. Funai, S. G. Drakos, J. Rutter, G. S. Ducker, Direct mitochondrial import of lactate supports resilient carbohydrate oxidation. bioRxiv (2024). doi:10.1101/2024.10.07.617073.

47. L. Chen, Y. Lin, X. Zhu, S. Zhuo, Z. Li, C. Guo, X. Ye, J. Chen, S. Wang, Y. Chen, MCT1-mediated Lactate Shuttle to Mitochondria Governs Macrophage Polarization and Modulates Glucose Homeostasis by Affecting beta Cells. Adv. Sci. 12, e14760 (2025).

48. W. Chai, W. Hao, J. Liu, Z. Han, S. Chang, L. Cheng, M. Sun, G. Yan, Z. Liu, Y. Liu, G. Zhang, L. Xing, H. Chen, P. Liu, Visualizing Cathepsin K-Cre Expression at the Single-Cell Level with GFP Reporters. JBMR Plus 7, e10706 (2023).

49. C. L. Winkeler, R. D. Kladney, L. B. Maggi, Jr., J. D. Weber, Cathepsin K-Cre causes unexpected germline deletion of genes in mice. PLoS One 7, e42005 (2012).

50. S. Morikawa, Y. Mabuchi, Y. Kubota, Y. Nagai, K. Niibe, E. Hiratsu, S. Suzuki, C. Miyauchi-Hara, N. Nagoshi, T. Sunabori, S. Shimmura, A. Miyawaki, T. Nakagawa, T. Suda, H. Okano, Y. Matsuzaki, Prospective identification, isolation, and systemic transplantation of multipotent mesenchymal stem cells in murine bone marrow. J. Exp. Med. 206, 2483–2496 (2009).

51. T. Sághy, P. Gupta, K. Horkeby, P. Henning, C. Ohlsson, C. Corciulo, M. K. Lagerquist, A. S. Chagin, C. Engdahl, Disrupted bone microenvironment and immune recovery following total body irradiation in a murine model. Cell Death Dis. 17, 122 (2026).

52. N. E. Putnam, L. E. Fulbright, J. M. Curry, C. A. Ford, J. R. Petronglo, A. S. Hendrix, J. E. Cassat, MyD88 and IL-1R signaling drive antibacterial immunity and osteoclast-driven bone loss during Staphylococcus aureus osteomyelitis. PLoS Pathog. 15, e1007744 (2019).

53. M. J. Campbell, C. Bustamante-Gomez, Q. Fu, K. E. Beenken, H. Reyes-Pardo, M. S. Smeltzer, C. A. O’Brien, RANKL-mediated osteoclast formation is required for bone loss in a murine model of Staphylococcus aureus osteomyelitis. Bone 187, 117181 (2024).

54. E. A. Masters, B. F. Ricciardi, K. L. de Mesy Bentley, T. F. Moriarty, E. M. Schwarz, G. Muthukrishnan, Skeletal infections: microbial pathogenesis, immunity and clinical management. Nat. Rev. Microbiol. 20, 385–400 (2022).

55. H. Li, S. Hong, J. Qian, Y. Zheng, J. Yang, Q. Yi, Cross talk between the bone and immune systems: osteoclasts function as antigen-presenting cells and activate CD4+ and CD8+ T cells. Blood 116, 210–217 (2010).

56. L. Ibáñez, G. Abou-Ezzi, T. Ciucci, V. Amiot, N. Belaïd, D. Obino, A. Mansour, M. Rouleau, A. Wakkach, C. Blin-Wakkach, Inflammatory Osteoclasts Prime TNFalpha-Producing CD4(+) T Cells and Express CX(3) CR1. J. Bone Miner. Res. 31, 1899–1908 (2016).

57. J. L. Krauss, P. M. Roper, A. Ballard, C. C. Shih, J. A. J. Fitzpatrick, J. E. Cassat, P. Y. Ng, N. J. Pavlos, D. J. Veis, Staphylococcus aureus Infects Osteoclasts and Replicates Intracellularly. mBio 10, e02447–19 (2019).

58. E. M. Palmieri, M. Gonzalez-Cotto, W. A. Baseler, L. C. Davies, B. Ghesquière, N. Maio, C. M. Rice, T. A. Rouault, T. Cassel, R. M. Higashi, A. N. Lane, T. W. M. Fan, D. A. Wink, D. W. McVicar, Nitric oxide orchestrates metabolic rewiring in M1 macrophages by targeting aconitase 2 and pyruvate dehydrogenase. Nat. Commun. 11, 698 (2020).

59. Z. Tan, N. Xie, S. Banerjee, H. Cui, M. Fu, V. J. Thannickal, G. Liu, The monocarboxylate transporter 4 is required for glycolytic reprogramming and inflammatory response in macrophages. J. Biol. Chem. 290, 46–55 (2015).

