## Supplemental Figures and tables for "Efferocytosis Remodels Osteoclast Metabolism and Enhances Antibacterial Defense"

Qixiu Yu^†,1^, Jie Zhang^†,1^, Ying Qu^†,1^, Yong Tang^1^, Yuyang Zhang^1^, Yewei Jia^1^,

Qijie Dai^1^, Zhoukai Huang^1^, Jie Shen^1^, Nathachit Limjunyawong^2,3^, Fei Luo^1^*, Ce Dou^1^*

^1^Department of Orthopedics, Southwest Hospital, Army Medical University, Chongqing, 400038, China

^2^Center of Research Excellence in Allergy & Immunology, Research Department, Faculty of Medicine Siriraj Hospital, Mahidol University, Bangkok 10700, Thailand.

^3^Department of Immunology, Faculty of Medicine Siriraj Hospital, Mahidol University, Bangkok 10700 Thailand

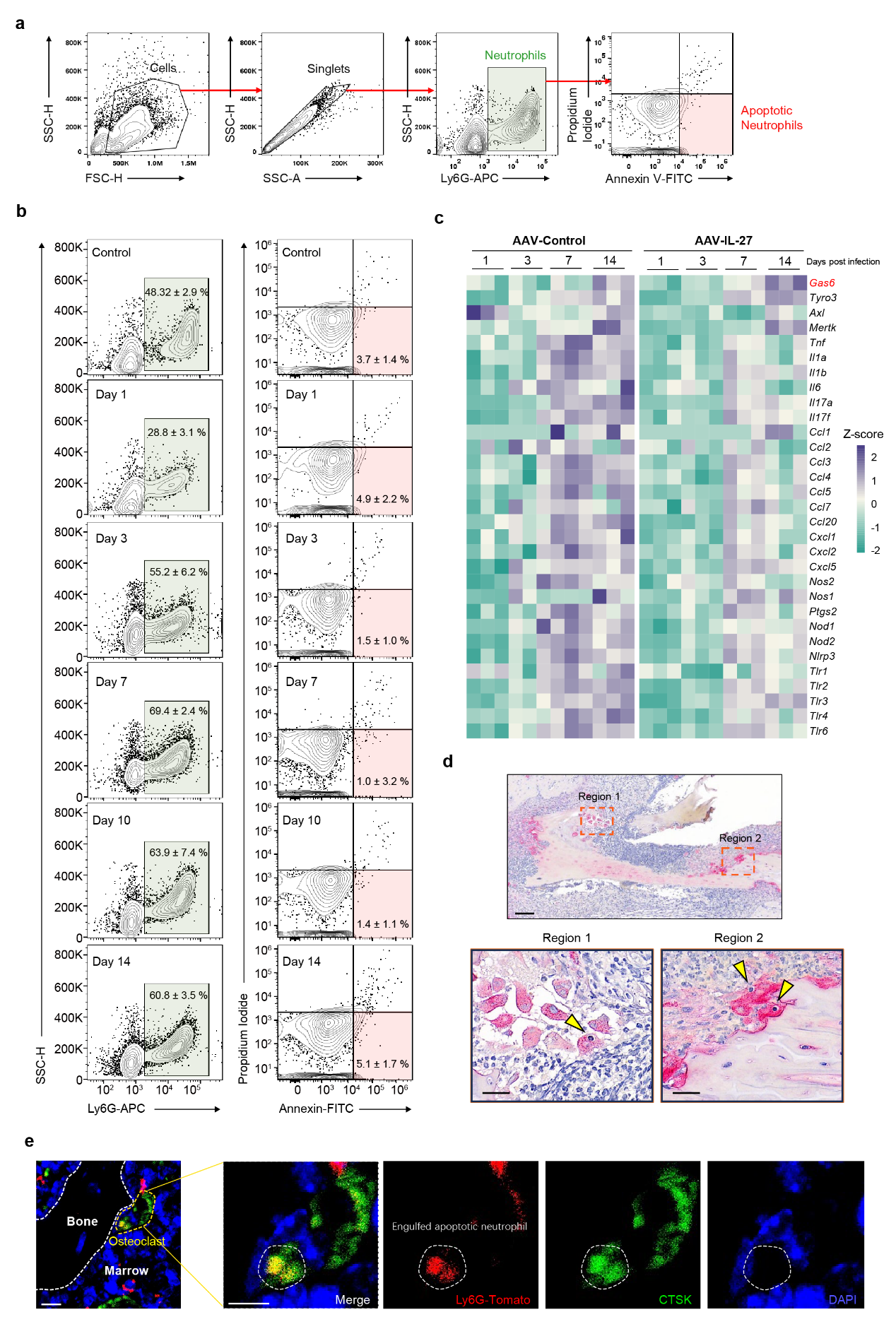
**Fig. S1. Apoptotic neutrophil accumulation and osteoclast efferocytosis in infected bone.**

**a,** Gating strategy used to identify bone marrow neutrophils (Ly6G⁺) and apoptotic neutrophils (Annexin V⁺/propidium iodide⁻) by flow cytometry.

**b,** Representative flow cytometry plots showing bone marrow neutrophil frequency (left) and apoptotic neutrophil frequency (right) in uninfected controls and at the indicated time points after *Staphylococcus aureus* infection.

**c,** Heatmap of transcripts encoding efferocytosis-associated molecules (*Gas6*, *Tyro3*, *Axl*, and *Mertk*), inflammatory cytokines, chemokines, and innate immune receptors in infected bone at the indicated time points, reanalyzed from a published dataset (GSE168896). n = 3 samples per treatment and time point; each column represents one infected-tibia RNA-seq sample.

**d,** Representative TRAP-stained femoral sections of the diaphyseal region after infection. Boxed areas (Regions 1 and 2) are shown at higher magnification. Arrows indicate osteoclasts undergoing efferocytosis. Scale bars, 100 µm (main images) and 25 µm (insets).

**e,** Additional representative bone sections from Ly6G-tdTomato reporter mice after infection. CTSK⁺ osteoclasts (green), tdTomato⁺ neutrophil-derived signals (red), and DAPI staining (blue) are shown. Arrows indicate neutrophil-derived signals within osteoclast boundaries. Scale bars, 10 µm (main images) and 5 µm (insets).

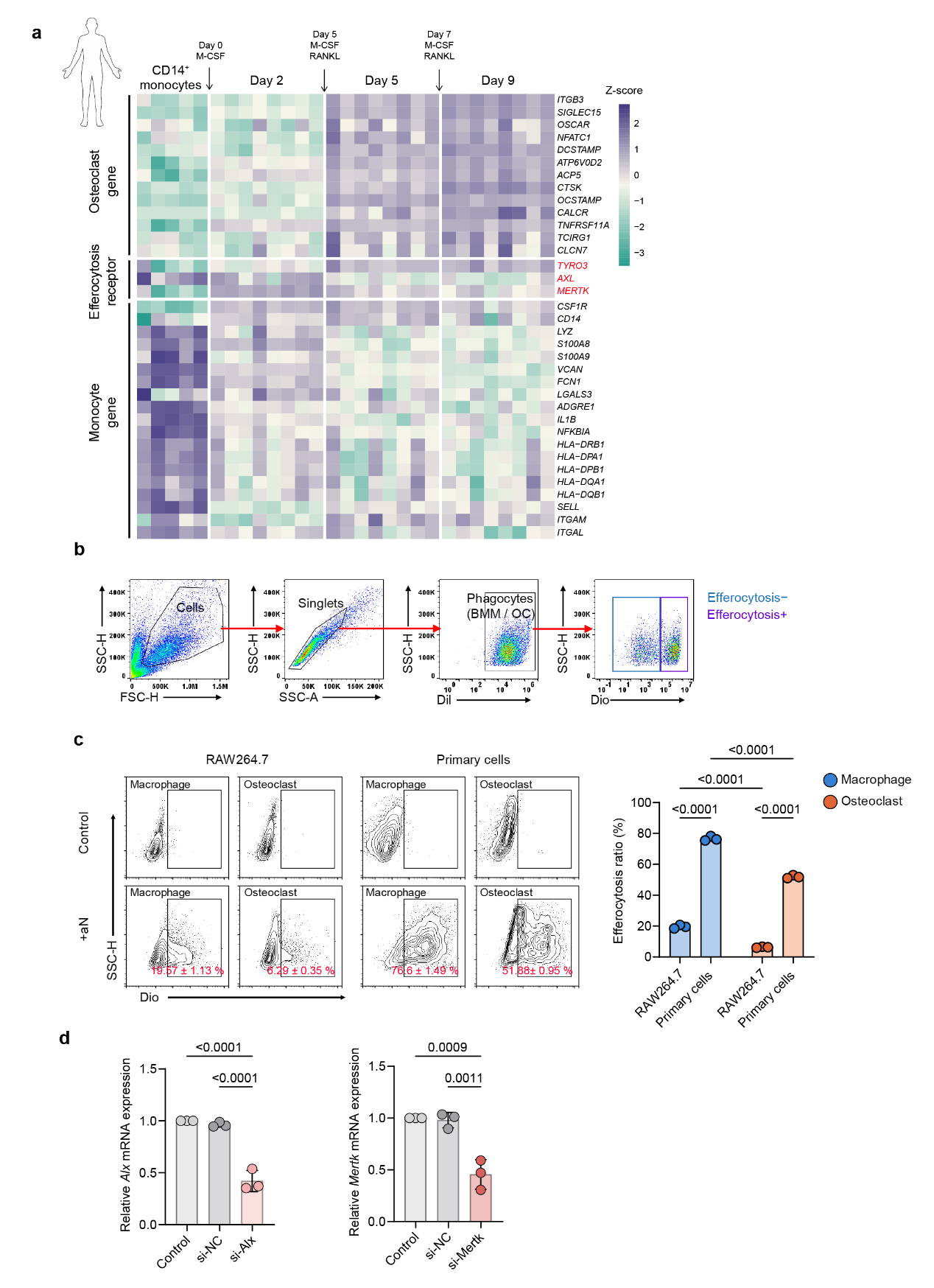
**Fig. S2. TAM receptor expression during human osteoclastogenesis and validation of efferocytosis assays.**

**a**, Heatmap showing expression of osteoclast genes, efferocytosis receptors (*TYRO3*, *AXL*, and *MERTK*) and monocyte genes during M-CSF- and RANKL-induced differentiation of human CD14⁺ monocytes into osteoclasts (GSE246769). Arrows indicate the timing of M-CSF and RANKL addition. n = 5 donor samples at day 0 and n = 8 at days 2, 5 and 9; each column represents one donor sample.

**b**, Gating strategy for flow cytometric quantification of efferocytosis. DiI-labeled phagocytes (BMMs or osteoclasts) were gated, and efferocytosis was defined by acquisition of the DiO signal from apoptotic neutrophils.

**c**, Representative flow cytometry plots and quantification of apoptotic neutrophil (aN) uptake by RAW264.7 cells and RAW264.7-derived osteoclasts, as well as by primary BMMs and osteoclasts.

**d**, RT–qPCR validation of siRNA-mediated knockdown of *Axl* (left) and *Mertk* (right) in BMMs. Expression levels were normalized to *Actb*.

Data are presented as mean ± s.d. Statistical significance was assessed using one-way ANOVA followed by Tukey’s multiple comparisons test (**c,d**). n = 3 independent biological samples per group (c,d); each point represents one independently generated primary-cell preparation or independently initiated RAW264.7 culture.

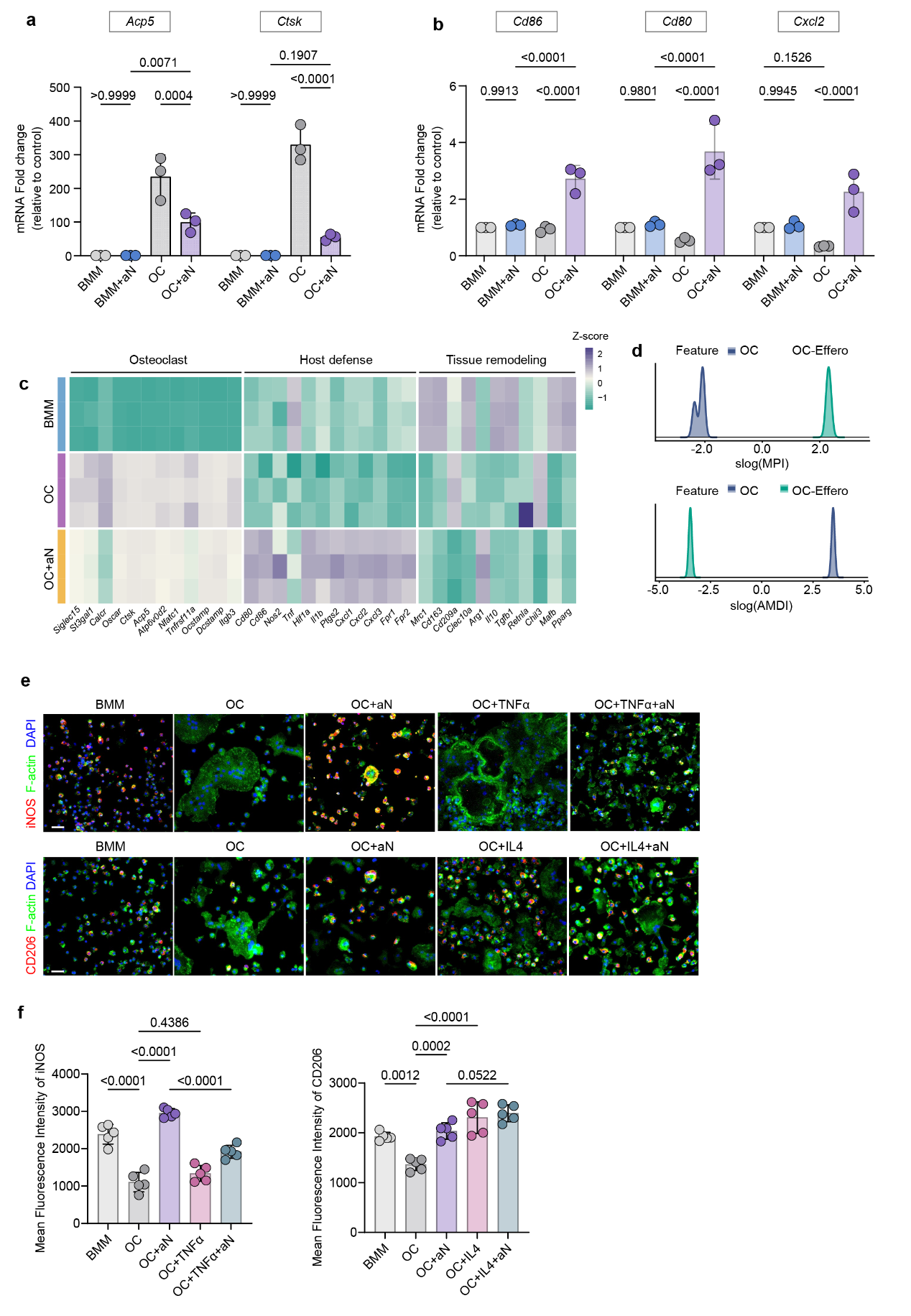
**Fig. S3. Transcriptional and functional consequences of apoptotic neutrophil efferocytosis in osteoclasts.**

**a,b**, RT–qPCR analysis of osteoclast-associated genes (*Acp5* and *Ctsk*) (**a**) and host defense-associated genes (*Cd86*, *Cd80*, and *Cxcl2*) (**b**) in BMMs and osteoclasts before and after apoptotic neutrophil (aN) efferocytosis. Expression levels were normalized to *Actb*.

**c**, Heatmap of osteoclast-associated, host defense, and tissue remodeling gene sets in BMMs, osteoclasts (OC), and efferocytic osteoclasts (OC+aN). n = 3 biological RNA-seq replicates per group; each row represents one biological sample.

**d**, MacSpectrum analysis of transcriptional similarities to macrophage activation states in osteoclast and efferocytic osteoclast cultures, showing the distribution of the macrophage polarization index (MPI, top) and the activation-induced macrophage differentiation index (AMDI, bottom). Analysis used n = 3 biological RNA-seq replicates per group.

**e,f**, Immunofluorescence analysis of iNOS (top) and CD206 (bottom) in BMMs and osteoclasts treated with TNF-α or IL-4 in the presence or absence of apoptotic neutrophils (**e**), with quantification of mean fluorescence intensity (**f**). iNOS and CD206 are shown in red, F-actin is shown in green, and nuclei are shown in blue (DAPI). Scale bars, 50 µm.

Data are presented as mean ± s.d. Statistical significance was assessed using one-way ANOVA followed by Tukey’s multiple comparisons test (**a,b,f**). n = 3 independent biological samples per group (a,b) and n = 5 (f); each point represents one biological sample.

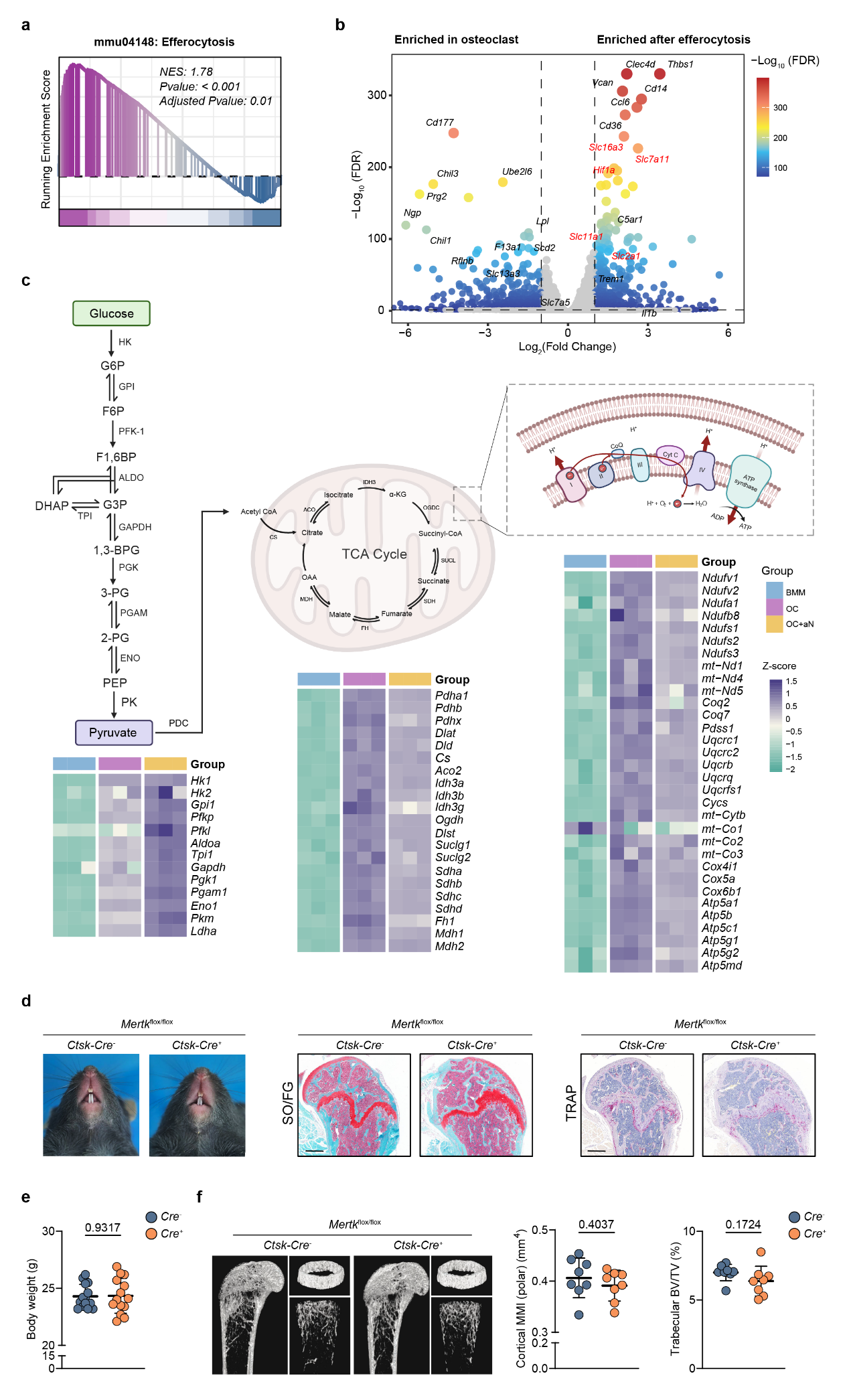
**Fig. S4. Metabolic features of efferocytic osteoclasts and baseline skeletal phenotypes of OC^(ΔMertk)^ mice.**

**a**, Gene set enrichment analysis of the efferocytosis pathway (mmu04148) in efferocytic osteoclasts relative to osteoclasts. NES, normalized enrichment score. Analysis used n = 3 biological RNA-seq replicates per group.

**b**, Volcano plot of differentially expressed genes between osteoclasts and efferocytic osteoclasts. Solute carrier transcripts and transcripts associated with glucose handling are highlighted in red. Each point represents one gene; the comparison used n = 3 biological RNA-seq replicates per group.

**c**, Schematic of glycolysis, the tricarboxylic acid (TCA) cycle, and oxidative phosphorylation, with heatmaps showing expression of the corresponding pathway genes. Related to Fig. 4a. n = 3 biological RNA-seq replicates per group; each heatmap column represents one sample.

**d**, Incisor eruption and representative Safranin O/Fast Green (SO/FG) and TRAP staining of femoral sections from OC^(ΔMertk)^ (*Ctsk*-Cre⁺; *Mertk*^flox/flox^) mice and Cre⁻ littermate controls. Scale bar, 500 µm.

**e**, Body weight of Cre⁻ and Cre⁺ littermates under steady-state conditions.

**f**, Representative micro-CT images of femurs from Cre⁻ and Cre⁺ littermates, with quantification of trabecular bone volume fraction (BV/TV) and cortical mean polar moment of inertia [MMI (polar)].

Data are presented as mean ± s.d. Statistical significance was assessed using two-tailed unpaired *t*-tests (**e,f**). n = 14 Cre− and 13 Cre+ mice (e) and n = 8 mice per genotype (f); each point represents one mouse.

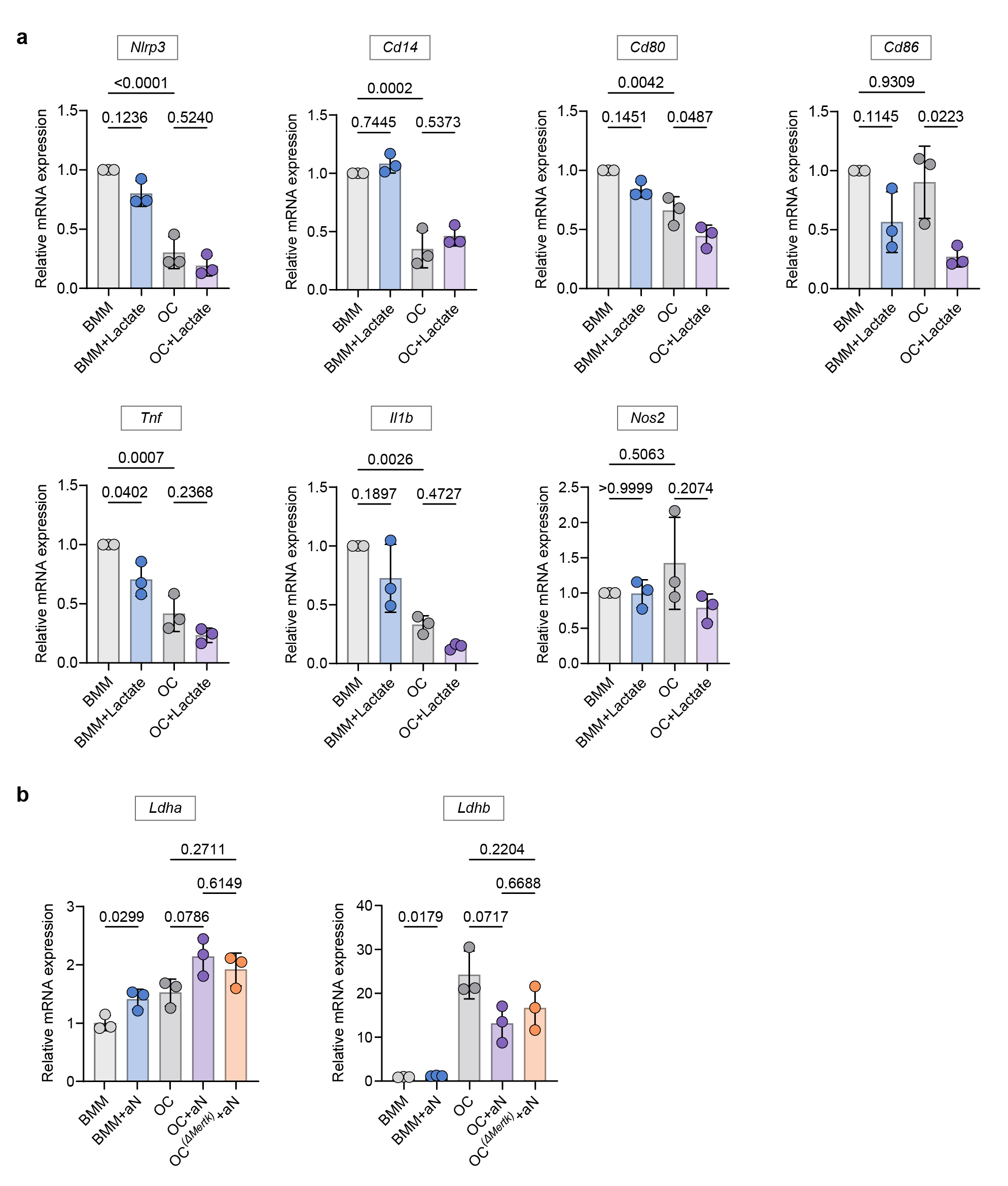
**Fig. S5. Analysis of immune effector and lactate metabolic gene expression.**

**a**, RT–qPCR analysis of immune effector genes (*Nlrp3*, *Cd14*, *Cd80*, *Cd86*, *Tnf*, *Il1b*, and *Nos2*) in BMMs and osteoclasts treated with or without sodium lactate. Expression levels were normalized to *Actb*.

**b**, RT–qPCR analysis of *Ldha* and *Ldhb* expression in BMMs and osteoclasts before and after apoptotic neutrophil efferocytosis. Expression levels were normalized to *Actb*.

Data are presented as mean ± s.d. Statistical significance was assessed using one-way ANOVA followed by Tukey’s multiple comparisons test (**a,b**). n = 3 independent biological samples per group (a,b); each point represents one biological sample.

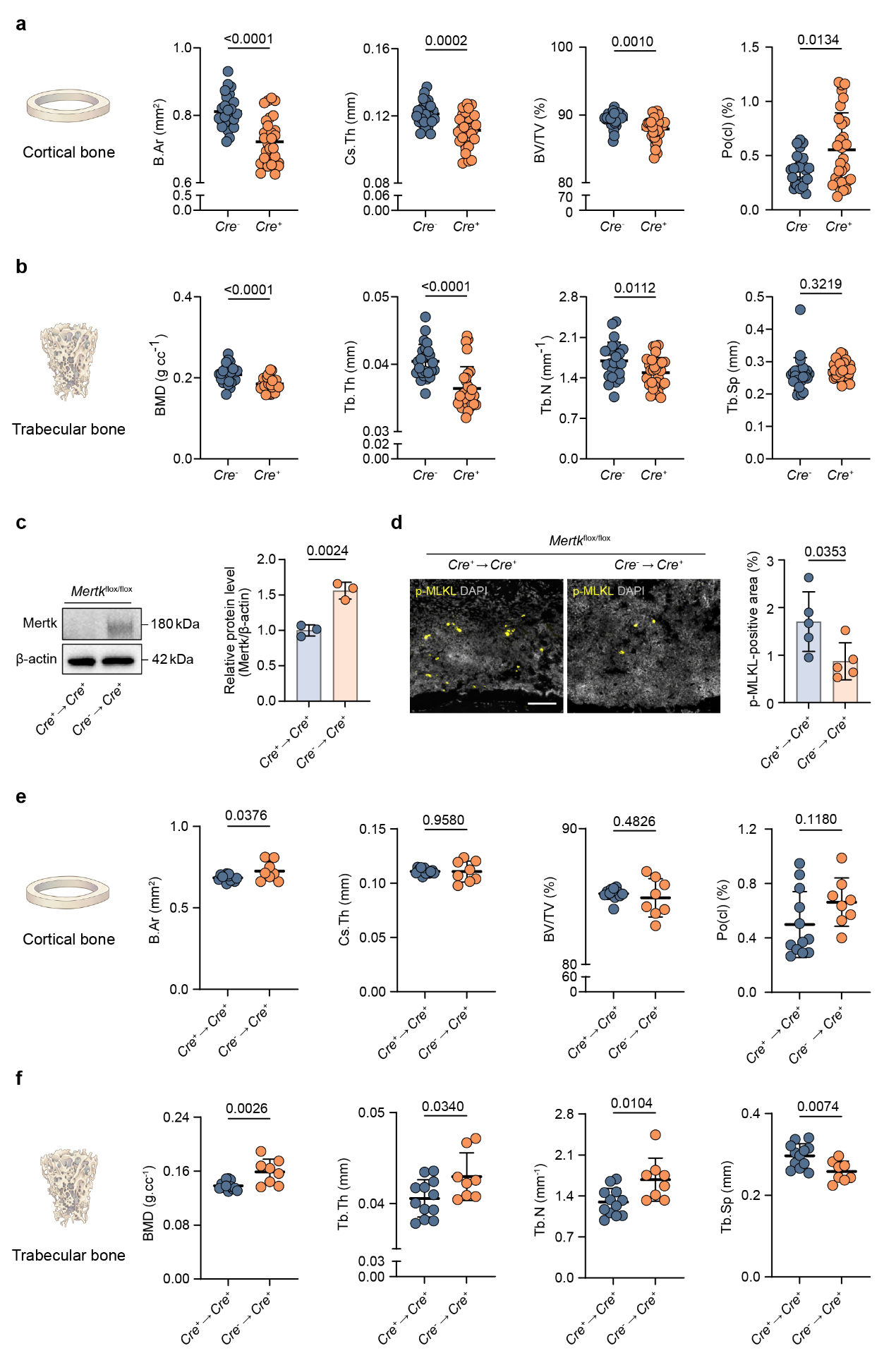
**Fig. S6. Additional characterization of osteoclast MERTK function during bone infection.**

**a,b**, Additional micro-CT parameters of femurs from OC^(ΔMertk)^ and littermate control mice at day 14 after *Staphylococcus aureus* infection. Cortical parameters (**a**): cross-sectional bone area (B.Ar), cross-sectional thickness (Cs.Th), bone volume fraction (BV/TV), and closed porosity (Po(cl)). Trabecular parameters (**b**): bone mineral density (BMD), trabecular thickness (Tb.Th), trabecular number (Tb.N), and trabecular separation (Tb.Sp).

**c**, Immunoblot analysis of MERTK expression in osteoclasts differentiated from bone marrow cells 6 weeks after BMT.

**d**, Histological analysis of bone tissues from BMT-reconstituted mice, with quantification of p-MLKL-positive areas (%). Scale bars, 100 µm.

**e,f**, Corresponding cortical (**e**) and trabecular (**f**) micro-CT parameters of femurs from BMT-reconstituted mice at day 14 after infection.

Data are presented as mean ± s.d. Statistical significance was assessed using two-tailed unpaired *t*-tests (**a,b,c,d,e,f**). n = 25 Cre− and 30 Cre+ mice (a,b); n = 3 recipient-derived cell preparations per group (c); n = 5 mice per group (d); n = 12 Cre+ BM→Cre+ and 8 Cre− BM→Cre+ mice (e,f). Each point represents one mouse, or one recipient-derived cell preparation in c.
